# *First-in-class* Transactivator-Free, Doxycycline-inducible IL-18-engineered CAR-T cells for relapsed/refractory B-cell lymphomas

**DOI:** 10.1101/2024.01.23.576842

**Authors:** Pedro Justicia-Lirio, María Tristán-Manzano, Noelia Maldonado-Pérez, Carmen Barbero-Jiménez, Marina Cortijo-Gutiérrez, Kristina Pavlovic, Francisco J Molina-Estevez, Pilar Muñoz, Ana Hinckley-Boned, Carmen Griñán-Lison, Saúl A Navarro-Marchal, Julia Muñoz-Ballester, Pedro A González-Sierra, Concha Herrera, Juan A Marchal, Francisco Martín

## Abstract

**Background:** Despite their success treating type B cancers, Chimeric Antigen Receptor (CAR) T cells still showed limited efficacy in certain lymphomas and solid tumors. Reinforcing conventional CAR-T cells to release cytokines can improve their efficacy but also increase safety concerns. Several strategies have been developed to regulate their secretion using minimal promoters that are controlled by chimeric proteins harboring transactivators. However, these chimeric proteins can disrupt the normal physiology of target cells.

**Methods:** Co-transduction with CAR19 and Lent-On-Plus-IL-18 LVs allowed for generating constitutive CAR/Dox-inducible IL-18 CAR-T cells that respond to ultra-low doses of doxycycline (iTRUCK19.18). iTRUCK19.18 were evaluated against an aggressive Burkitt lymphoma model *in vitro* and *in vivo*, against primary B-cell tumors and against a CD19-engineered pancreatic tumor model. Patient-derived iTRUCK19.18 cells were also generated.

**Results:** iTRUCK19.18 controlled IL-18 release through a dual mechanism dependent on doxycycline and T cell activation, thereby enhancing the safety profile. IL-18 release increased the activation state/proinflammatory profile of T cells in a doxycycline-dependent manner without altering cellular fitness, which was translated into an increased CAR-T cell antitumor activity against aggressive hematologic and solid tumor models. In a clinically relevant context, we generated patient-derived iTRUCK19.18 cells able to significantly increase elimination of primary B cells tumors under doxycycline. Furthermore, IL-18-releasing iTRUCK19.18 polarized pro-tumoral M2 macrophages towards an antitumoral phenotype (M1), suggesting the ability to modulate the tumor microenvironment.

**Conclusion:** We have generated the first transactivator-free inducible TRUCKs from healthy donors and B-cell neoplasms patients. iTRUCK19-18 exhibit dual safety control mechanisms for IL-18 secretion and improved antitumoral activity against type-B neoplasms. Inducible IL-18 secretion not only enhanced T cell potency but could also change the tumor microenvironment to a more antitumoral state.

## Background

CD19-redirected Chimeric Antigen Receptor (CAR)-T cell therapy has provided long-lasting clinical responses treating relapsed and/or refractory (R/R) B-cell neoplasms (1). The high complete remission rate achieved in patients unresponsive to multiple lines of treatment has led the FDA and EMA to approve six CAR-T-based Advanced Therapy Medicinal Products (ATMPs) to date.

Despite outstanding results in the treatment of different CD19+ hematologic cancers, around 30-50% of patients relapse after αCD19-CAR-T infusion (2). CAR-T therapy has shown limited therapeutic efficacy in other hematological malignancies such as chronic lymphocytic leukemia (CLL) (3), and very few reports have shown efficacy on solid tumors (4). Several factors appear to be influencing the loss of CAR-T efficacy: 1) CAR-derived tonic signaling, that accelerates functional exhaustion and toxicity of CAR-T cells (5); 2) impaired long-term CAR-T cell persistence (6); 3) restricted trafficking of CAR-T cells into the tumor (7); and 4) a highly immunosuppressive tumor microenvironment (TME).

One of the possible solutions to improve the therapeutic efficacy of CAR-T involves the fine-tuning of TRUCKs (T cells redirected for antigen-unrestricted cytokine-initiated killing), CAR-T cells engineered to release transgenic proteins that boost their antitumor capacity. Generally, TRUCKs co-express interleukins such as IL-7, IL-12, IL-15 or IL-18. These key cytokines exert an immunomodulatory role in reversing the pro-tumor environment of the TME into an antitumor milieu, promoting a coordinated immune cell attack on the tumor (8). However, the expression of these cytokines in long-lived T cells requires tight regulation to avoid a plethora of potential unwanted side effects. Up to date, most TRUCKs designs use endogenously regulated NFAT (Nuclear Factor of Activated T cell)-based promoters to express cytokines, to generate tumor-specific T cells that secrete the selected cytokine only after T cell activation (9). However, several studies suggest the inability of the NFAT promoter to efficiently control cytokine expression inside the tumor (10, 11). This likely arises from the presence of multiple signals beyond just CAR- or TCR-binding to its target that can initiate NFAT activation. These signals include G Protein-Coupled Receptors (GPCR) signaling (involved in T cell migration and activation) (12), proinflammatory cytokines (IL-2, IL-4, IL-6 and TNFα) (13) and viral infections (14). Therefore, regulating T cell potency by NFAT-driven promoters remains challenging. Alsaieedi *et al.* discovered that while the expression of IL-12 was necessary to observe antitumor effectiveness, the introduction of NFAT-IL12 transgenic T cells into a syngeneic murine model of B16F10 melanoma led to lethality (11). In the same direction and in a clinical context, Zhang *et al.* observed severe toxicity in melanoma patients treated with autologous tumor infiltrating lymphocytes (TILs) genetically engineered to express IL-12 under NFAT promoter (10).

As an alternative to endogenously regulated NFAT-driven promoters, several groups are developing exogenously inducible promoters (15), such as the one based on the bacterial TetO operon. In this direction Alsaieedi *et al*. showed that, while NFAT-driven IL-12 engineered T cells induce lethality (see above), Tet-On engineered T cells were safe in the absence of doxycycline (Dox) and that temporal induction of IL-12 inhibits the growth of B16F10 melanoma tumors. This data evidence the potency of Dox-inducible Tet-On systems as tools to generate smart ATMPs that can be controlled externally by clinicians. However, most Tet-On systems require a transactivator (a chimeric protein composed of the bacterial Tet repressor (TetR) and the activating domain of the viral protein 16 (VP16) of herpes simplex virus type 1) to achieve inducibility. These transactivators showed multiple side effects on gene-modified cells due to transcription factor sequestering or by binding to pseudo TetO sites (16–19). Importantly, Smith *et al*. observed a depletion of antigen-experienced T cells in transactivator (rtTA)-transgenic mice, evidencing the potential difficulties of using these systems to generate clinical-grade inducible T cells. To tackle these problems, our group has previously developed insulated (20), transactivator-free, Tet-On LVs (Lent-On-Plus or LOP) (21, 22) that tightly regulate transgene expression in a variety of primary human cells, including T cells, without altering physiology and using ultra-low doses of Dox (23).

Previous studies have shown that constitutive expression of IL-18 by CAR-T cells significantly enhances the antitumor activity of CAR-T cells. IL-18 is a cytokine of the IL-1 family constitutively produced by activated macrophages, dendritic and epithelial cells. It directly stimulates interferon gamma (IFNγ) secretion and other inflammatory cytokines and chemokines, exhibiting pleiotropic effects on the entire immune system via Th1 immune response (24). IL-18 enhances the cytotoxic activity of T cells and natural killer (NK) cells by upregulation of FasL (25), polarization of pro-tumorigenic M2 to antitumor M1 macrophages (26), and by acting in synergy with other cytokines (27). Recently, it was reported that IL-18-secreting CAR-T cells showed superior antitumor activity via helper effect of CD4+ CAR-T cells for the augmentation of CD8+ CAR-T cells (28). Avanzi *et al*. demonstrated that IL-18 CAR-T cells were able to eliminate liquid and solid tumors in syngeneic murine models (29). Taking all these data, the University of Pennsylvania’s team is currently running a clinical trial (NCT04684563, phase 1) co-expressing IL-18 on αCD19-CAR-T cells to evaluate the maximum safe dose.

As for most cytokines, unregulated release of IL-18 can lead to potential safety issues, since a continuous delivery of IL-18 promotes constant (and non-tissue-specific) IFNγ secretion, creating a permanent environment of acute inflammation that might lead to toxicity (30), autoimmune disorders (31) as well as IFNγ-independent toxicities (32). Therefore, a controlled/regulated release of IL-18 is highly desirable. In this regard, Chmielewski *et al*. generated CAR-T cells able to release IL-18 ‘on demand’, using a NFAT promoter. The authors showed that their engineered CAR-T cells released IL-18 in a CAR-dependent fashion and increased the antitumor effect compared with standard CAR-T cells (26). However, as outlined above, several studies suggest the inability of the NFAT promoter to efficiently control cytokine expression *in vivo* (10, 11).

As mentioned before, Dox-based inducible expression systems have emerged as interesting alternatives to control the expression of cytokines with the limitation of high Dox requirements and the presence of highly toxic transactivators. Here we describe *first-in-class* αCD19-CAR-T cells (iTRUCK19.18) engineered to release IL-18 under ultra-low (subtherapeutic) Dox doses and in the absence of transactivators. iTRUCK19.18 controlled IL-18 expression both *in vitro* and *in vivo*, allowing the control of T cell potency and polarizing pro-tumoral M2 macrophages towards an antitumoral phenotype (M1) in a dox-dependent manner. This effect was translated into an increased CAR-T cell antitumor activity against an aggressive hematologic and an engineered-CD19+ PDAC model. In a clinically relevant context, we also generated patient-derived iTRUCK19.18 and observed that the Dox-dependent release of IL-18 improved the eradication of primary B cell tumors.

## Methods

### Cell lines

HEK-293T (human embryonic kidney derived cells, ATCC CRL-11268) and MIA-PaCa2 (human pancreatic adenocarcinoma cells, ATCC CRL-1420) cell lines were cultured with DMEM (Dulbecco’s Modified Eagle Medium) (Biowest) supplemented with 10% fetal bovine serum (FBS) (Gibco) and 1% penicillin/streptomycin (P/S) (Gibco). Jurkat (acute T-cell leukemia, ATCC TIB-152) and Namalwa (Burkitt’s lymphoma cells, ATCC CRL-1432) cell lines were grown in RPMI-1640 (Roswell Park Memorial Institute) (Biowest) supplemented with 10% FBS and 1% P/S. Cell lines were routinely tested for mycoplasma.

### Isolation and culture of primary T cells

Peripheral blood samples from healthy donors and patients were provided by the Hematology and Hemotherapy Unit of the Reina Sofía University Hospital (Córdoba) and Virgen de las Nieves University Hospital (Granada) under informed consent, following the guidelines of the ethics committee and in accordance with Spanish regulations (RD-L 9/2014). T cells from healthy donors were obtained from peripheral blood mononuclear cells (PBMCs). Blood was diluted 1/2-1/4 in PBS (Biowest) and PBMCs were isolated using Ficoll gradient centrifugation (Cytiva) at 400g for 20 minutes without brake nor acceleration. The mononuclear lymphocyte layer was collected and washed with PBS. Cells were cultured at 2×10^6^ cells/ml in TexMACS medium (Miltenyi Biotec) supplemented with 10 ng/ml of IL-7 and IL-15 (Miltenyi Biotec) and 1% P/S (Biowest) in at 37°C and 5% CO_2_.

Patients’ PBMCs were obtained from 5-8 ml of blood from patients diagnosed with marginal zone lymphoma (MZL) or chronic lymphocytic leukemia (CLL) prior to treatment. PBMCs were isolated as described before and cultured at 2×10^6^ cells/ml with TexMACS medium supplemented with 1% P/S, 5% human AB serum (Biowest) and 40 IU/ml of IL-2 (Miltenyi Biotec). 24 hours later, TransAct (1:100) was added for 6 days prior transduction. Expansion after transduction was performed with TexMACS supplemented with 10 ng/ml of IL-7 and IL-15, 1% P/S and 5% human AB serum.

### Lentiviral vectors

EF1α-A3B1-19BBz (CAR19, ARI-0001) plasmid was kindly provided by Dr. Manel Juan and Dr. Maria Castella from Hospital Clínic (Barcelona, Spain). CIL18ETIS2 (LOP18) plasmid was generated by designing and incorporating hIL18 sequence (RefSeq Transcript ID: NM_001386420.1, synthesized by ATG:biosynthetics) flanked by AscI/SbfI sites into the CELETIS2(23) plasmid, replacing the eGFP-2A-Nluc region.

### Lentiviral vectors production and titration

HEK-293T cells were co-transfected with the transfer plasmid, plasmid pCMVDR8.91 and plasmid pMD.G as described in (22) using polyethylenimine (PEI) (Alfa Aesar). Viral supernatants were collected 48 and 72h post transfection and concentrated 100x by ultracentrifugation (90000g, 4°C, 2 hours) and functional titre was determined in Jurkat cells as described (33).

### Generation of iTRUCK19.18

Primary T cells were activated with T Cell TransAct (Miltenyi Biotec) and 48 hours later co-transduced by a mixture of CAR19 LVs and LOP18 LVs. Briefly, LVs were mixed and cells were added for spinoculation (800g, 32°C, 1h). 5 hours later, cells were washed and plated at a density of 1×10^6^ with TexMACS medium (Miltenyi Biotech) supplemented with 10 ng/ml of IL-7 and IL-15 (Miltenyi Biotech).

### Flow cytometry

CAR expression was analyzed using a primary goat IgG1 antibody that binds to the murine Fab conjugated with biotin (AB_2632441,1:100, Jackson Immunoresearch) and Streptavidin-APC (17-4317-82, 1:330, eBioscience). Briefly, an anti-Fab antibody was added for 40 minutes. After washing, Streptavidin-APC was added for 30 minutes. After 15 minutes in the presence of Streptavidin, surface antibodies were added.

For the immunophenotyping of primary T cells, the following monoclonal antibodies were used: CD45RA-FITC (HI-100, 1:200), CD62L-PE-Cy7 (DREG56, 1:200), CD3-PerCP-Cy5.5 (OKT3, 1:200), CD4-eFluor450 (RPA-T4, 1:100), PD1-APC (MIH4, 1:50), LAG-3-PE (3DS223H, 1:100), and TIM-3-eFluor780 (F38-2E2, 1:50), all from eBioscience (ThermoFisher). For measuring AICD markers, the following monoclonal antibodies were used: anti-Human CD253 (TRAIL)-PE (RIK-2, 1:100, BD Biosciences), anti-Hu CD95 (APO-1/Fas)-APC (DX2, 1:200, eBiosciences), AnnexinV/7ADD (eBiosciences) and Fas Ligand Ms Anti-Hu mAb-FITC (SB93a, 1:200, Life Technologies). For the characterization of primary macrophages, CD206-FITC (19.2, 1:100), CD11c-PE (3.9, 1:50) and CD14-PerCP-Cy5.5 (61D3, 1:200) from eBiosciences (ThermoFisher) were used.

Intracellular staining was performed using the Fix & Perm kit (Nordic MUbio), following manufacture’s recommendations. Anti-hIL-18 Propeptide-PE (74801, 1:20, R&D systems), pCD3ζ-PE (Tyr142, 3ZBR4S, 1:100, eBioscience), anti-Hu IFNgamma-FITC (4S.B3, 1:100, eBioscience), anti-Hu Granzyme B-eFluor 450 (N4TL33, 1:100, eBioscience), anti-Hu IL-2-PE-Cyanine7 (MQ1-17H12, 1:100, eBioscience), anti-Hu TNFalpha-APC (Mab11, 1:100, eBioscience) and anti-Hu Perforin-FITC (d69, 1:100, eBiosciences) were used for different assays.

Mice samples were obtained by mechanical disruption (bone marrow, spleen and liver) or after a Percoll (GE Healthcare) gradient (brain). Fcγ receptors were blocked using murine αCD16/CD32 (ThermoFisher), human FcR Blocking (Miltenyi Biotec), and 5% mouse serum (Sigma Aldrich). All stainings were performed in darkness, on ice, and washes were performed with FACS buffer (PBS + 3% BSA + 2 mM EDTA) at 400g for 5 minutes, unless otherwise indicated. Cytometers used for acquisition were FACS Canto II and FACS Verse (BD Biosciences), performing exclusion by singlets, as well as live/dead cells using 4’,6-diamidino-2-phenylindole (DAPI, Thermo Fisher). In the case of intracellular stainings, cell viability was determined using Ghost Dye Violet 510 (TONBO biosciences). Absolute cell quantification was performed using CountBright Absolute Counting Beads (ThermoFisher). Data analysis was performed using FlowJo V10 software (TreeStar).

### IL-18 secretion assay

For the study of secreted hIL-18, we used HEKBlue IL-18 cells (Invitrogen) that allow the detection of bioactive hIL-18 (10 pg/ml – 1 ng/ml). Briefly, 3×10^5^ T cells were plated per condition in 150 μl of TexMACS medium (Miltenyi Biotec) without cytokines. Cells were activated with TransAct (1:150, Miltenyi Biotec) or PBS (negative activation control) for 24h. Supernatants were collected and frozen and then analyzed following manufacturer’s instructions.

### Polarization assay

PBMCs from healthy donors were thawed in a 96-well plate at a density of 2×10^6^ cells/ml. After 24 hours, suspended cells (mostly T cells) were separated and cultured with TexMACS (Miltenyi Biotec) supplemented with IL-7 and IL-15 (Miltenyi Biotec), while the adherent cells (monocytes/macrophages) were cultured with RPMI-1640 supplemented with 10% FBS (Biowest). Macrophages were supplemented with either 1) 50 ng/ml GM-CSF (PeproTech) (for polarization to an M1 phenotype) or 2) 50 ng/ml M-CSF (PeproTech) (for polarization to an M2 phenotype). 96 hours later, macrophages were supplemented with either 1) 10 ng/ml of E. coli lipopolysaccharide (LPS, Sigma Aldrich) for final polarization to M1 or 2) 20 ng/ml of IL-4 (PeproTech) for final polarization to M2. Flow cytometry analysis was performed after 6 days to confirm macrophage polarization.

### Cytotoxicity assays

<u>Namalwa cells:</u> briefly, 7.5×10^4^ Namalwa GFP-Nluc cells were co-cultured with T cells in duplicate in 96-well plates (ThermoFisher), maintaining an effector-to-target (E:T) ratio of 1:10 (calculated based on CAR+ cells), in TexMACS medium without supplementation. Tumor re-stimulations were performed by adding the same number of tumor cells as initially present.

<u>Primary tumors:</u> 5×10^4^ primary CD19+ tumor cells isolated from peripheral blood of untreated patients with MZL and CLL were co-cultured with T cells from healthy donors or from the same patient, transduced with CAR and LOP18, at three different E:T ratios: 1:1, 1:2, and 1:5 (calculated based on the percentage of CAR+ cells), in duplicate in 96-well plates. Cytotoxicity reading was performed 13 hours after co-culturing using flow cytometry.

<u>MIA PaCa2 CD19+ cells:</u> 7.5×10^3^ MIA PaCa2 target cells expressing 100% GFP-Nluc and 70% CD19 were seeded in duplicate in 96-well plates (ThermoFisher) the day before adding T cells. Cells were cultured in complete DMEM (Biowest) (+10% FBS, +1% P/S) (Biowest). Next day, target cells were incubated with T cells at an E:T ratio of 1:2 (calculated based on CAR+ cells) in non-supplemented TexMACS medium for 48 hours. Tumor re-stimulations were performed by adding the same number of tumor cells as initially present.

Specific lysis was calculated using the following formula: Specific lysis = 1 - (%CD19+ cells in CAR condition / %CD19+ in NT condition) * 100; formula adapted from (34).

### Cytokines quantification

Cytokines quantification has been performed using the MACSPlex Cytokine 12 Kit (Miltenyi Biotec), following the manufacturer’s instructions. Log2 (score) was calculated as follows: Log2(secreted cytokine in iTRUCK condition/secreted cytokine in CAR condition), using the mean between donors.

### Animal models

All mice were handed according to EU European (2010/63/UE) and local animal wellness regulations (RD1386/2018, RD53/2013), previous revision and approval by the local Ethics Committee. To generate the Burkitt’s lymphoma murine model, 0.3×10^6^ Namalwa GFP-Nluc cells were intravenously inoculated into 10 to 12-week-old NOD/scid-IL-2Rnull (NSG) mice. Three or six days later, CAR-T cells were intravenously injected. In some cases, tumor re-inoculations were performed.

Mice were humanely euthanized by cervical dislocation when they exhibited a high bioluminescence signal, a weight loss of >20% of their initial weight or showed clear signs of pain or xenogeneic graft-versus-host-disease (xenoGVHD).

### Bioluminescence analysis

Bioluminescence analysis was performed as described (23). Briefly, furimazine (NanoGlo, Promega) was administered via intraperitoneal just before acquiring the image using the IVIS Spectrum analyzer (Caliper, Perkin Elmer). Images were analyzed using Living Image 3.2 (Perkin Elmer) or AURA Imaging 4.0.7 (Spectral Instruments Imaging).

### Data analysis

Statistical analyses were performed using GraphPad Prism 9 (Dotmatics). Data is expressed as mean ± SEM. Each n represents an independent donor; each N represents a mouse. Statistical test was indicated in the corresponding figure caption.

## Results

### Generation of transactivator-free, IL-18-inducible αCD19 CAR-T cells (iTRUCK19.18)

As recently described by our group, LOP LVs can tightly regulate transgene expression in primary T cells *in vitro* and *in vivo* (23). Based on this data, we decided to generate CAR-T cells that express IL-18 in an inducible manner (iTRUCK19.18) using the LOP system. To achieve this, we co-transduced primary T cells with CAR19 LVs (multiplicity of infection, MOI=3) (allowing constitutive expression of a 4-1BB αCD19 CAR endowed with the A3B1 scFv clone) (35) (**Fig. 1a**, top) and LOP18 (MOI=5) (for dox-inducible expression of IL-18 using the Lent-On-Plus LV) (**Fig. 1a**, bottom), resulting in a heterogeneous population of cells including CAR+ IL-18+, CAR+ IL18-, CAR-IL18+, and CAR-IL18-cells. Co-transduction with two LVs seems to reduce CAR expression compared to transduction with single CAR19 LVs (**Fig. 1b**), probably due to LVs dilution and free VSV-G protein competing for free receptors. Importantly, 7-9 days post-transduction, iTRUCK19.18 showed minimal intracellular pro IL-18 expression in the absence of dox (**Fig. 1c**, left, blue histogram and **Fig. 1c**, right, blue dots) and up to 44.1% in its presence (**Fig. 1c**, left, green histogram and **Fig. 1c**, right, green dots).

**Fig. 1.**
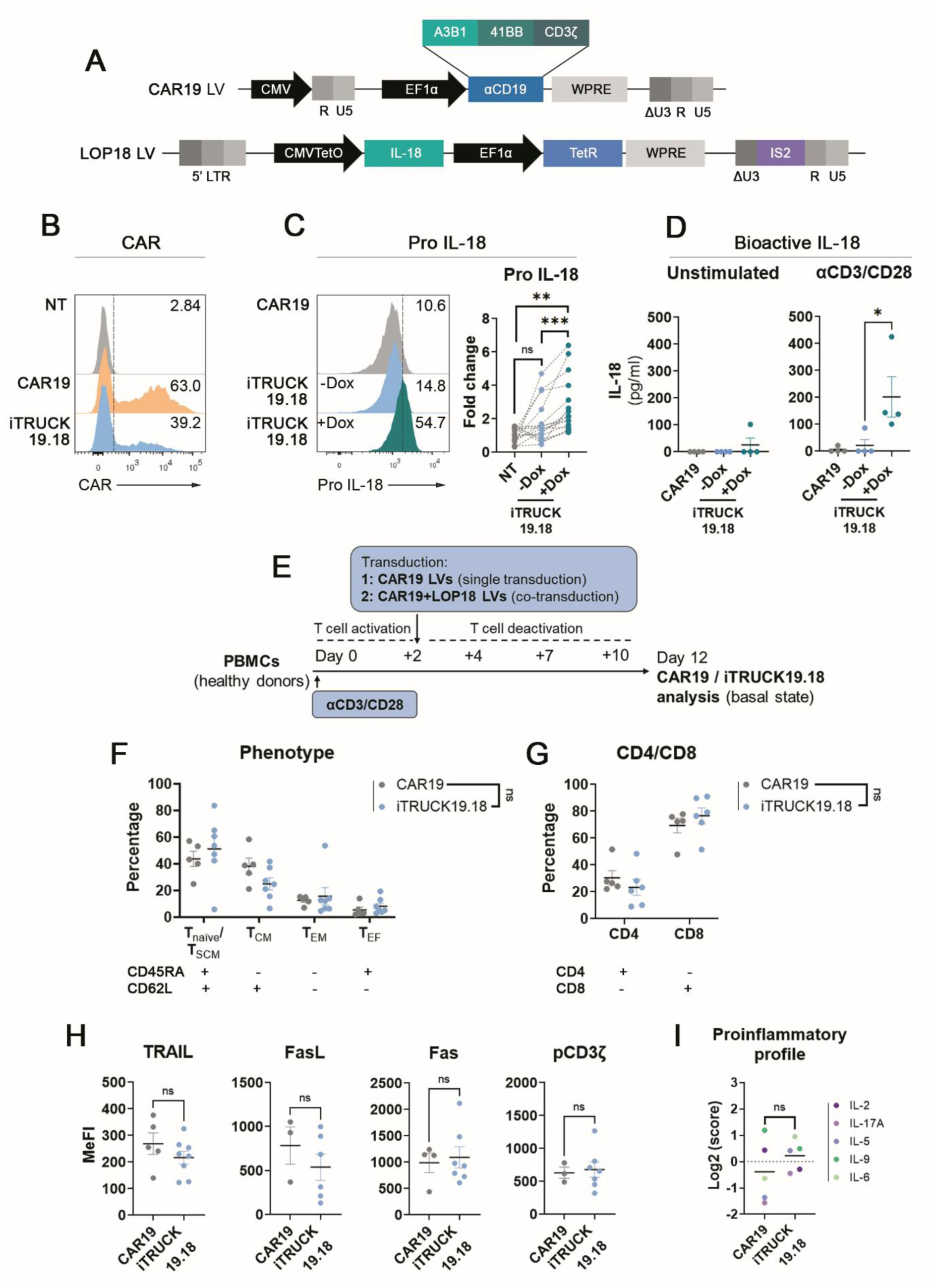
Generation and characterization of inducible IL-18-producing CAR-T cells (iTRUCK19.18). **a** Schematic representation of CAR19 LV encoding for EF1α-A3B1-41BB-CD3ζ (top) and for the transactivator-free, dox inducible LOP LVs expressing pro-IL-18 (bottom). **b** Representative histograms of CAR expression in non-transduced cells (NT, top), in CAR19 (middle) and in iTRUCK19.18 -Dox (bottom). **c** Representative histograms of pro IL-18 expression in CAR19 cells (top) and iTRUCK19.18 cells in the absence (middle) or presence (bottom) of 50 ng/ml Dox. Right: Fold change of IL-18 expression from iTRUCK19.18 cells relative to basal expression of non-transduced cells (NT: n=17; -Dox: n=15; +Dox: n=17) (right). **d** Detection of bioactive IL-18 secreted by iTRUCK19.18 without activation (left) and activating with TransAct (αCD3/CD28) (right) (n=4). **e** Experimental procedure for CAR19 and iTRUCK19.18 cells generation (single or co-transduction with LVs) and their analysis in resting conditions after 10 days in the absence of stimuli, mimicking the endpoint of the manufacturing process (infusion product, without Dox). **f** Phenotype of CAR19 and iTRUCK19.18 cells in the absence of Dox of the infusion product (ARI-0001: n=5; iTRUCK19.18: n=7). **g** CD4/CD8 ratio of CAR19 and iTRUCK19.18 cells in the absence of Dox of the infusion product (ARI-0001: n=5; iTRUCK19.18: n=7). **h** Expression of AICD markers (TRAIL, FasL, Fas) and phosphorylation of CD3z in CAR19 vs iTRUCK19.18 cells (-Dox): TRAIL (CAR19: n=5; iTRUCK19.18: n=8), FasL (CAR19: n=3; iTRUCK19.18: n=6), Fas (CAR19: n=4; iTRUCK19.18: n=7), pCD3ζ (CAR19: n=3; iTRUCK19.18: n=7). **i** Index of proinflammatory cytokines-related secretion by CAR19 vs iTRUCK19.18 cells at basal state (n=2). *p<0.05, **p<0.01, ***p<0.001 (two-tailed paired t test).

Once we confirmed that iTRUCK19.18 cells induced pro IL-18 in a dox-dependent manner, our subsequent aim was to verify the accurate processing and secretion of the cytokine. Physiologically, IL-18 is synthesized as a pro-peptide mainly by activated macrophages, dendritic and epithelial cells. Activation signal or tissue damage triggers the pro-caspase 1 processing into functional caspase 1 by the inflammasome complex, converting pro IL-18 into mature IL-18, which is secreted (36). We therefore analyzed if transgenic IL-18 expressed by iTRUCK19.18 cells follows a similar process despite being expressed in a non-natural context. Our findings are consistent with this mechanism, as iTRUCK19.18 cells necessitates T cell stimulation in conjunction with the presence of Dox to secrete bioactive IL-18 (**Fig. 1d**). This observation aligns with the natural processing of IL-18, endowing these cells with a dual switch mechanism involving both T cell-activation and Dox exposure. This configuration enhances their safety profile.

With the aim of characterizing the generated product, we conducted a comparative analysis of iTRUCK19.18 cells and CAR19 cells to decipher whether co-transduction affected production and T cell fitness compared to the generation of conventional CAR-T cells. We analyzed the immunophenotype, CD4/CD8 cell ratio, activation-induced cell death (AICD) markers, signaling through CD3ζ phosphorylation (pCD3ζ), and the proinflammatory profile by measuring the secretion of five proinflammatory cytokines at the end of the production process (12 days, in basal state) (**Fig. 1e**). We found no significant differences between iTRUCK19.18 cells and CAR19 cells in any of the parameters analyzed (**Fig. 1f-i**), indicating the feasibility of generating iTRUCK19.18 by co-transduction with CAR19 and LOP18 LVs.

### Dox addition to iTRUCK19.18 cells enhanced their activation capacity without compromising T cell exhaustion and phenotype

Once it was confirmed that the production process of iTRUCK19.18 cells was feasible, we analyzed the effect of IL-18 production on T cells (**Fig. 2a**). IL-18 plays a pivotal role in the activation of T cells, so initially, we evaluated the effect of IL-18 production at basal level. To evaluate the effect on AICD, we analyzed TRAIL, FasL, and Fas expression in iTRUCK19.18 cells under both dox-induced and non-induced conditions. Upon dox exposure, we observed an upregulation of TRAIL (**Fig. 2b**, first graph), a tendency in FasL (**Fig. 2b**, second graph) but no difference was observed in Fas expression (**Fig. 2b**, third graph). Interestingly, Dox administration led to a significant increase in the expression of phospho-CD3ζ (**Fig. 2b**, fourth graph) (representative dot-plots on **Fig. S2a**). However, despite the notable changes in AICD and tonic signaling, we did not observe any differences in apoptosis in the presence of Dox (**Fig. 2b**, right graph).

**Fig. 2.**
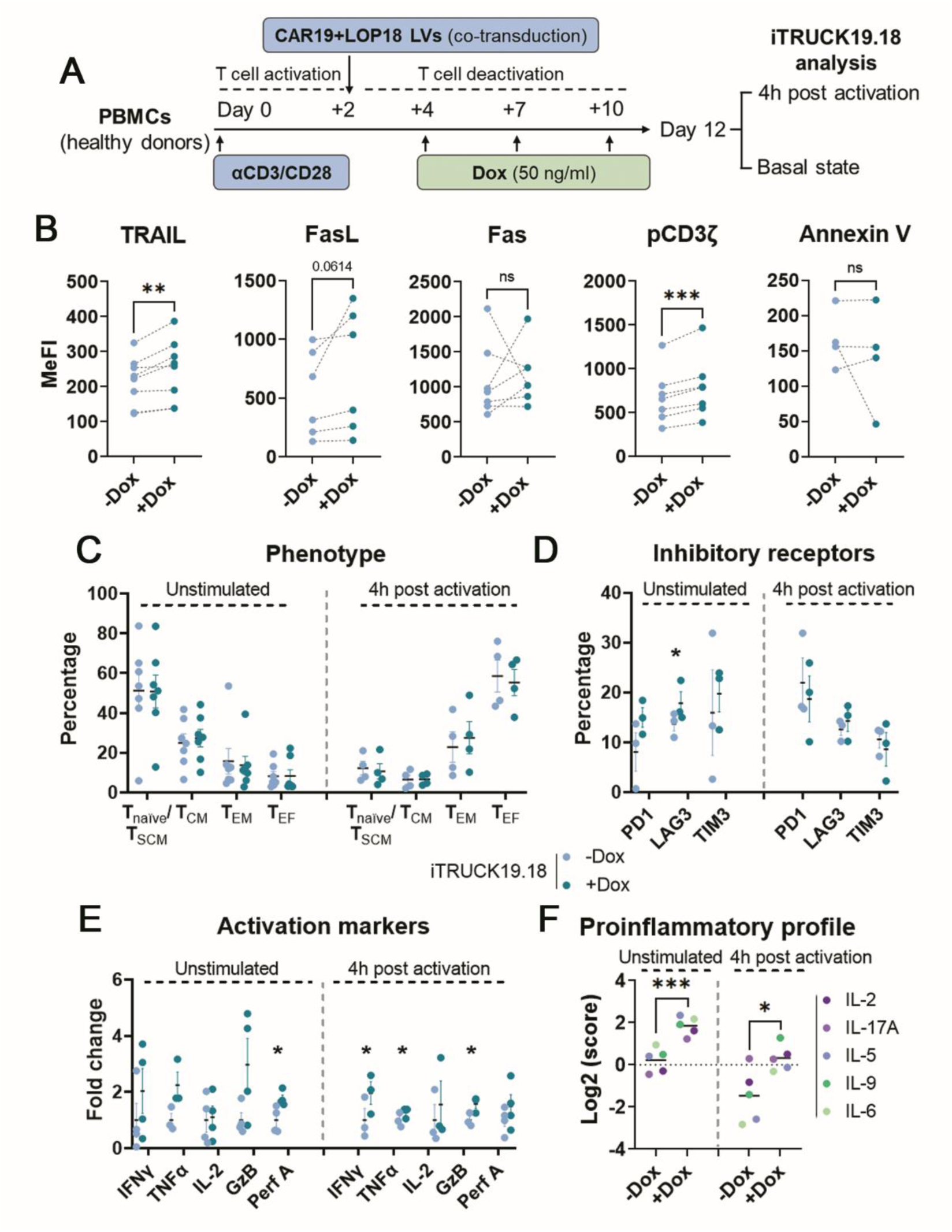
Characterization of iTRUCK19.18 cells in absence and presence of dox. **a** Experimental design for iTRUCK19.18 cells generation and analysis at basal state (simulating the endpoint of the manufacturing process) or after 4h of activation with αCD3/CD28. **b** Expression of AICD markers (TRAIL, FasL, Fas), phosphorylation of CD3z and apoptosis marker Annexin V (from left to right) in iTRUCK19.18 cells at basal state in the absence (light blue) or presence of 50 ng/ml Dox (dark blue): TRAIL (n=8), FasL (n=6), Fas (n=7), pCD3ζ (n=7) and Annexin V (n=4). **c** Phenotype of iTRUCK19.18 cells with (dark blue) and without Dox (light blue) at resting conditions (left) (n=7) or after 4h of stimulation (right) (n=4). **d** Percentage of positive cells for exhaustion markers PD1, LAG3, TIM3 of iTRUCK19.18 cells at basal state and after stimulation in the absence or presence of 50 ng/ml Dox (n=4). **e** Fold change of the activation markers IFNγ, TNFα, IL-2, Granzyme B and Perforin A on iTRUCK19.18 cells without (light blue) or with Dox (dark blue) at resting state (left) or after activation (right) (n=4). **f** Proinflammatory index of iTRUCK19.18 cells with and without Dox at basal state (n=2). Each dot represents a different cytokine as indicated in the right panel. *p<0.05, **p<0.01, ***p<0.001 (one-tailed paired t test).

Next, we aimed to conduct a more comprehensive characterization of the fitness of iTRUCK19.18 cells, not only at the basal level but also after stimulation. At a basal level, IL-18 production did not alter the distribution of phenotypic subpopulations (**Fig. 2c**) or the ratio of CD4+/CD8+ T cells (**Fig. S2b**). Additionally, the expression of inhibitory receptors PD1, LAG3, and TIM3 on T cells remained unaffected, except for a significant increase in the expression of LAG3 in the basal state. However, upon activation, we observed that this increase in LAG3 expression didn’t persist (**Fig. 2d**). This suggests that the production of this cytokine does not accelerate T cell exhaustion. As expected due to the release of IL-18, iTRUCK19.18 cells exposed to Dox increased the expression of several activation markers. Following cellular activation, we observed a significant upregulation in the expression of IFNγ, TNFα, and Granzyme B, along with a trend towards higher expression of IL-2 and Perforin A. Interestingly, Perforin A was significantly increased under basal conditions in the presence of Dox (**Fig. 2e**). The increased activation state of iTRUCK19.18 cells in response to dox is consistent with the augmentation of their proinflammatory profile, as evidenced by the upregulated secretion of IL-2, IL-17A, IL-5, IL-9, and IL-6 (**Fig. 2f** and **Fig. S2c**). Of note, we also observed a trend in the increase of IL-4 and IL-10 (**Fig. S2d**), underlining the dual role of the cytokine depending on the context.

It is important to notice that no significant differences were found in single transduced CAR19 cells (without LOP18) by the use of Dox in any of the markers analyzed either at basal state or after activation (**Fig. S3a-e**), indicating that the observed effects are exclusively due to the production of IL-18 and not by the Dox *per se*. Considering all these data, we conclude that Dox addition increased the activation of iTRUCK19.18 cells while retaining the exhaustion state without significant phenotypic alterations.

### Doxycycline regulates the antitumoral activity of iTRUCK19.18 cells in a Burkitt lymphoma model *in vitro* and *in vivo*

Once demonstrated that the secretion of functional IL-18 by iTRUCK19.18 cells can be controlled by dox, maintaining an appropriate phenotype, we analyzed whether we can also control their antitumoral activity. For this purpose, we first co-cultured iTRUCK19.18 and CAR19 cells with Namalwa cells, a Burkitt lymphoma cell model, using serial tumor stimulations (**Fig. 3a**). The results demonstrated that during the third tumor encounter, IL-18-releasing iTRUCK19.18 cells exhibited a significant enhanced antitumoral activity compared to cells without Dox and standard CAR19 cells. Furthermore, even the dox-free condition displayed greater antitumoral action than CAR19 cells, suggesting that the initial secretion of IL-18 during the initial days post-transduction (the system requires 6-10 days post-transduction to achieve tight regulation) is having a positive effect on their fitness/antitumoral activity (**Fig. 3b**).

**Fig. 3.**
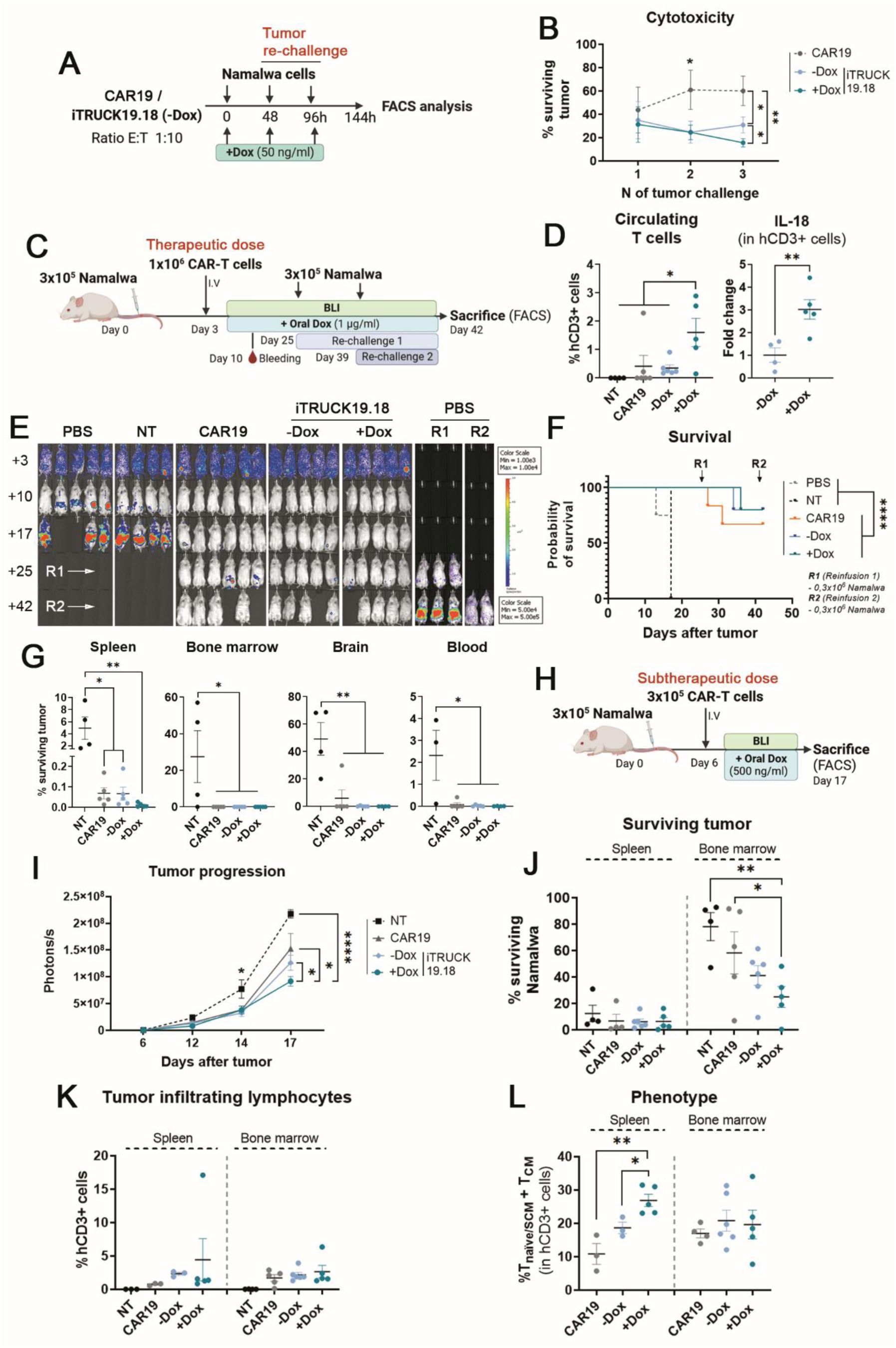
*In vitro* and *in vivo* evaluation of iTRUCK19.18 cells against B-cell lymphoma model. **a** Diagram representing the *in vitro* cytotoxicity assay. Briefly, CAR19 and iTRUCKs19.18 (always cultured in the absence of Dox) were co-cultured with tumor cells (Namalwa) every 48h for 3 tumor challenges at ratio 1:10 CAR-T:tumor cell. Dox was added at the moment of the co-culture and dose was refreshed in every challenge. **b** Percentage of surviving Namalwa cells after serial tumor encounters with CAR19 or iTRUCK19.18 cells without and with Dox (n=4). **c** In vivo experimental procedure to evaluate iTRUCK19.18 at a therapeutic dose (1×106 CAR-T cells/mouse). Dox (1000 ng/ml) was added to drinking water after infusion into mice and refreshed twice a week. Two more tumor challenges with Namalwa cells were infused at day 25 and day 42, respectively. **d** Proportion of circulating human T cells (left) and relative expression of IL-18 (right) by T cells obtained from blood 7 days after infusion of iTRUCK19.18 cells. e Bioluminescence images of tumor progression in mice treated with PBS, NT, CAR19 and iTRUCK19.18 without and with Dox. As control of rechallenges 1 and 2 (R1 and R2), novel mice were also infused with PBS at days +25 and +39. **f** Survival graph of mice treated with PBS, NT, CAR19, iTRUCK19.18 without and with Dox. **g** Percentage of viable tumor cells in different organs (spleen, bone marrow, brain and blood, from left to right) of mice treated with NT, CAR19 and iTRUCK19.18 without and with Dox, at final point. (PBS: N=5; NT: N=4; CAR19: N=6; iTRUCK19.18 -Dox: N=5; iTRUCK19.18 +Dox: N=5). **H** Diagram representing the infusion of a subtherapeutic dose (3×105 CAR-T cells) into mice 6 days post-tumor. Dox (500 ng/ml) was added to drinking water after the infusion of the CAR-T cells into mice and was refreshed twice a week. **i** Tumor progression determined by bioluminescence (photons/s) of the different experimental groups (NT, CAR19, iTRUCK19.18-Dox, and +Dox). **j** Percentage of surviving tumor cells in the spleen (left) and bone marrow (right) of the mice from the different experimental groups. **k** Percentage of tumor infiltrated T cells (hCD3+) in spleen (left) and bone marrow (right) of mice at the time of sacrifice. **L** Proportion of T_naive/SCM+TCM_ cells in the spleen (left) bone marrow (right) of mice at endpoint. (NT: N=4; CAR19: N=5; iTRUCK19.18 -Dox: N=6; iTRUCK19.18 +Dox: N=5). *p<0.05, **p<0.01, ****p<0.0001 (one-tailed paired t test for B; Log-Rank test for F; one-tailed unpaired t test for D, G, I, J and L).

Next, we assessed the *in vivo* efficacy of iTRUCK19.18 cells. For this purpose, 0.3×10^6^ Namalwa GFP-Nluc cells were infused into immunocompromised NSG mice and tumor progression was monitored after intravenous administration of 1×10^6^ iTRUCK19.18 cells (in the presence or absence of Dox), CAR19 cells, non-transduced T cells (NT) and PBS (**Fig. 3c**). After seven days of CAR-T cells administration, we observed a significant increase in human circulating T cells in the blood of mice treated with iTRUCK19.18 only under oral Dox supplementation (**Fig. 3d, left**). Furthermore, we observed that human T cells from mice treated with Dox significantly induced IL-18 expression (**Fig. 3d, right**), demonstrating that oral administration of Dox allows *in vivo* induction of IL-18.

In this context (with a high quantity of CAR-T and a low tumor burden), no differences in antitumoral response were observed between the different groups, as all mice treated with CAR19/iTRUCK19.18 completely eradicated lymphoma even after two additional tumor re-infusions (re-challenge 1 and 2, R1 and R2) (**Fig. 3e, f**). Interestingly, the analysis of tumor cells infiltration in different tissues showed that iTRUCK19.18-treated mice completely cleared tumor cells in all tissues in the absence or presence of dox, while 1/5 CAR19 cells-treated mouse presented brain metastasis (**Fig. 3g**). Unfortunately, we stopped the experiment on day 42 due to the development of xenoGVHD.

Since the high CAR-T cell dose prevented from seeing differences in antitumor potency, we performed a new *in vivo* experiment where we infused a subtherapeutic dose (0.3×10^6^ CAR-T cells/mouse) to mice under a higher tumor burden (allowing tumor expansion 6 days) (**Fig. 3h**). In this new scenario, we did observe how the addition of Dox increased the antitumor potency of iTRUCK19.18 cells compared with iTRUCK19.18 -Dox, CAR19 and NT-treated mice (**Fig. 3i**). On day 17 we stopped the experiment (due to systemic progression of the lymphoma), and we analyzed the presence of surviving tumor cells, and the phenotype of the infiltrated T cells in the spleen and bone marrow. No significant differences were found in terms of infiltrated tumor cells (surviving Namalwa) among the groups in spleen (**Fig. 3j**, **left**). However, mice treated with iTRUCK19.18 cells in the presence of Dox significantly reduced tumor cell invasion in bone marrow (**Fig. 3j**, **right**). No significant differences were observed in the infiltration of human T cells in the spleen and bone marrow among the different groups of mice treated with CAR-T cells (**Fig. 3k**). Furthermore, in the spleen of mice treated with iTRUCK19.18 +Dox, the infiltrated T cells exhibited a less differentiated/more memory phenotype (CD45RA+ CD62L+ and CD45RA-CD62L+) (**Fig. 3l, left**), which might have contributed to the enhanced antitumoral efficacy. However, no discernible differences were observed in the bone marrow concerning the phenotype of mice infused with CAR-T cells (**Fig. 3l**, **right**).

### Healthy donor and patient-derived iTRUCK19.18 cells show an increased antitumor potency against primary B-type tumors in the presence of Dox

After confirming that the production of IL-18 by CAR-T cells enhanced the antitumoral effect against a Burkitt lymphoma model, we wanted to validate the use of iTRUCK19.18 cells in a clinically relevant setting. We first selected primary tumors from three patients with B-cell neoplasms (patient #1 diagnosed with marginal zone lymphoma (MZL) and patient #2 and #3 with chronic lymphocytic leukemia (CLL), expressing heterogeneous levels of CD19 (**Fig. 4a**) and investigated the antitumoral efficacy of iTRUCK19.18 cells (derived from healthy donor T cells) in the presence or absence of Dox. Interestingly, and as observed for the lymphoma cell line, iTRUCK19.18 cells efficiently eliminated P#1 and P#2 both primary tumors in a Dox-dependent manner under the most restrictive effector:target ratio (**Fig. 4b**).

**Fig. 4.**
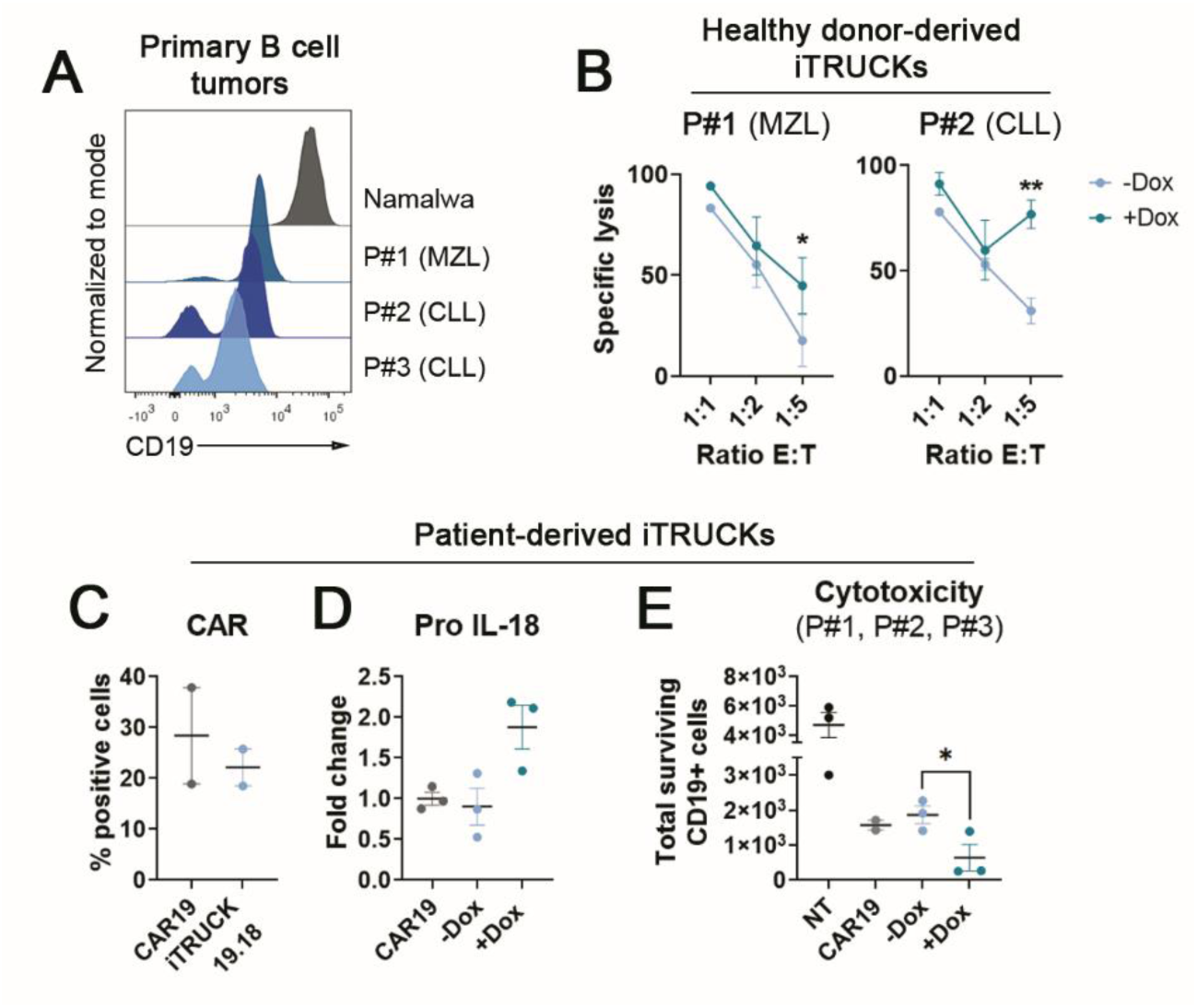
Characterization and lytic capacity of healthy donor and patient-derived iTRUCK19.18 cells against primary B tumors. **a** Histogram showing CD19 expression of primary tumor samples derived MZL (patient 1, leukemic mantle cell lymphoma) and CLL (patient 2 and 3, chronic lymphocytic leukemia) and Namalwa cells (from Burkitt’s lymphoma). **b** Specific lysis (compared to NT intrinsic lysis) by healthy donor-derived iTRUCK19.18 cells (generated from three healthy donor’s PBMCs) with (50 ng/ml) and without Dox against different proportions of P#1 (left) and P#2 (right) tumor samples (n=3, each patient’s sample were cocultured with iTRUCKs generated from three different donors). **c** Percentage of CAR+ cells of patient-derived CAR19 and iTRUCK19.18 cells without and with Dox (50 ng/ml) treatment (n=2, P#2, P#3). Briefly, CD3+ cells were enriched after αCD3/CD28 stimulation for 5 days and then, those cells were transduced with CAR19 and iTRUCK19.18 LVs. CAR expression was determined 5 days after transduction. **d** Fold change of pro IL-18 expression of patient’s derived CAR19 and iTRUCK19.18 after 48h in the presence of 50 ng/ml Dox (n=3). **e** Number of surviving CD19+ tumor cells following encounter with patient-derived iTRUCK19.18 cells at an E:T ratio of 1:5 after 13 hours of co-culture (NT, iTRUCK19.18 -Dox, and +Dox: n=3; CAR19: n=2). *p<0.05, **p<0.01 (two-tailed paired t test).

Next, we aimed to verify the feasibility of generating iTRUCK19.18 cells from patient’s T cells (**Fig. S4a**). Similar to the observations made when modifying T cells from healthy donors, the co-transduction showed a tendency to reduce CAR expression (**Fig. 4c**). As expected, the addition of Dox resulted in the induction of pro IL-18 (**Fig. 4d**). We further assessed a series of baseline markers in patient-derived iTRUCK19.18 cells and found results consistent with our previous observations (**Fig. S4b**). Interestingly, after serial encounters, we observed a trend towards decreased TOX expression after dox treatment (**Fig. S4c**), a marker associated with terminal exhaustion program in T cells. Finally, in an autologous setting, we corroborated that patient-derived iTRUCK19.18 cells exhibited enhanced efficacy in eliminating the same-patient tumor B cells only when IL-18 is induced by Dox (**Fig. 4e**). These compelling findings underscore the therapeutic potential of iTRUCK19.18 cells in treating B-type hematological neoplasms.

### Dox treatment on iTRUCK19.18 cells increases their antitumoral potency against metastatic CD19+ PDAC model

Based on the results obtained from applying iTRUCK19.18 cells to hematologic cancer models, we aimed to investigate potential future applications of LOP18 LVs to enhance CAR-T therapy against solid tumors. In this line, we use iTRUCK19.18 against a metastatic pancreatic ductal adenocarcinoma (PDAC) model engineered to express CD19 (MIA-PaCa2 cells 70% CD19) previously developed by our group (37). We analyzed the anti-tumor efficacy of iTRUCK19.18 cells *in vitro,* after serial tumoral challenges (**Fig. 5a**). The results revealed a remarkable increase in the antitumoral potency of iTRUCK19.18 cells with Dox compared to those without Dox, in every encounter analyzed (**Fig. 5b, S5a**). This enhanced effectiveness was linked to the maintenance of a less differentiated phenotype (T_naïve/SCM_+T_CM_) observed in the IL-18-producing cells starting from the second encounter (**Fig. 5c, S5b**). Additionally, we found no differences in PD1 or TIM3 exhaustion markers regardless of the Dox addition (**Fig. 5d, S5c**), so we could also confirm that IL-18-releasing iTRUCK19.18 cells not only don’t accelerate T cell exhaustion, but also retain T cells in a memory phenotype that increase their antitumoral potency.

**Fig. 5.**
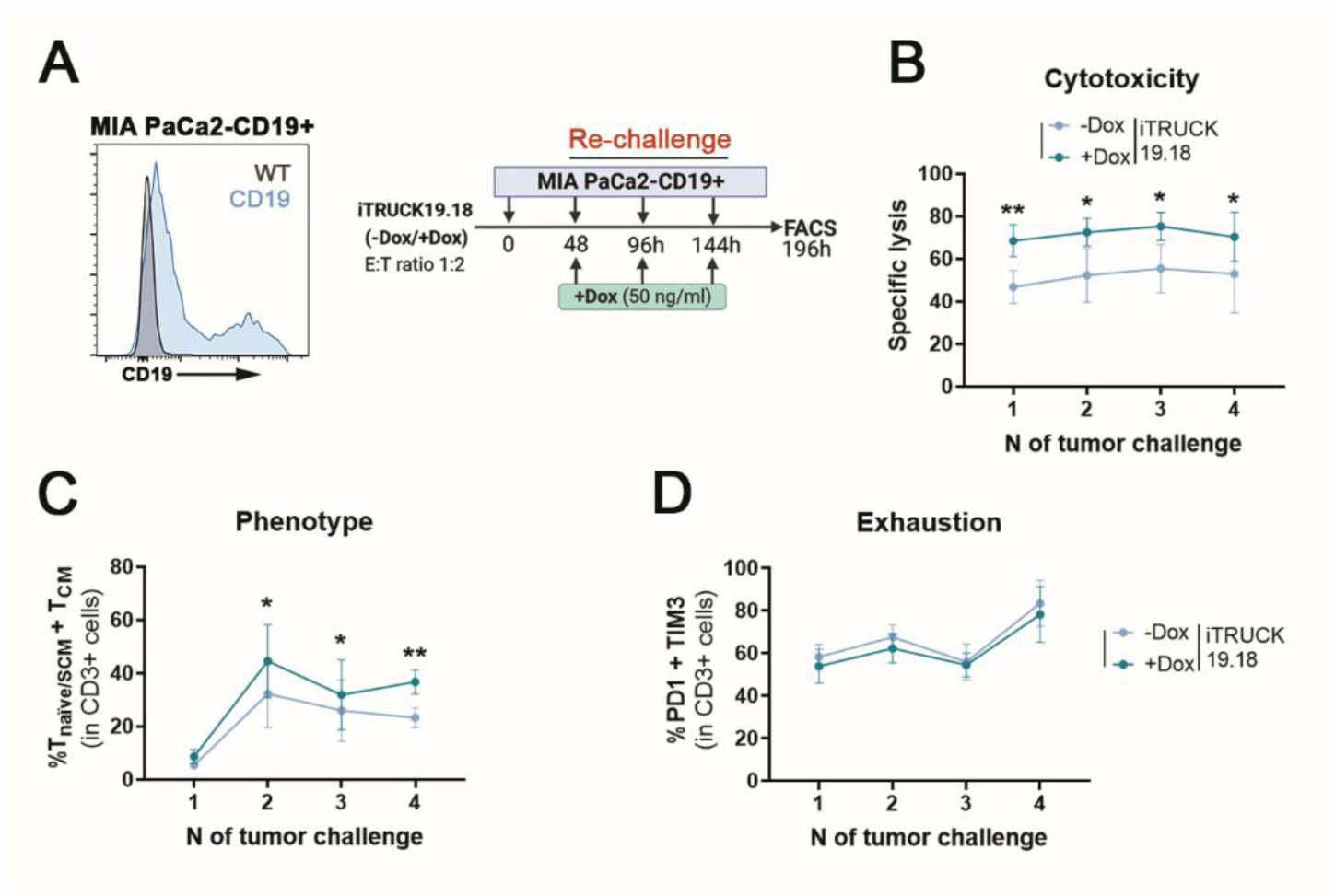
*In vitro* efficacy of iTRUCK19.18 cells against CD19+ pancreatic ductal adenocarcinoma tumor model. **a** Experimental procedure of the in vitro cytotoxicity assay with an artificial model of PDAC cells, MIA-PaCa2-CD19+, modified to express heterogeneous levels of CD19. Left: Histogram of the expression of CD19 of MIA-PaCa2 WT (grey) or CD19+ (blue), as target of CAR19 and iTRUCK 19.18. Right: Briefly, iTRUCK19.18 cells in the absence or presence (50 ng/ml) of Dox were co-cultured at ratio 1:2 effector: target with MIA-PaCa2-CD19+ cells. Tumor re-challenges and FACS analysis were performed every 48h. **b** Specific lysis over 4 tumor encounters of iTRUCK19.18 -Dox (light blue) and +Dox (dark blue) compared to NT (n=5). **c** Proportion of T_naïve/SCM_ and T_CM_ from -Dox (light blue) and +Dox (dark blue) iTRUCK19.18 after 48h of every tumor encounter (n=5). **d** Proportion of PD1+ TIM3+ cells analyzed at every tumor encounter (n=5). *p<0.05, **p<0.01 (two-tailed paired t test).

These findings collectively support the therapeutic potential of the LOP system in CAR-T cells for treating different tumor contexts by exogenously controlling IL-18 through Dox administration.

### IL-18 induction allows the control of the polarization of pro-tumoral macrophages towards an antitumoral phenotype

As mentioned before, the TME constitutes a major barrier to the clinical efficacy of CAR-T therapy not only against solid tumors, but also against hematological malignancies, resulting in suboptimal outcomes. Within the TME, tumor associated macrophages (TAMs) play a pivotal role, characterized by an M2 phenotype that exerts potent immunosuppressive effects, thereby constraining the functionality of CAR-T cells.

It has been demonstrated that the expression of IL-18 by CAR-T cells reduced the quantity of M2 macrophages in murine models (26). Consequently, we sought to investigate whether controlling the expression of IL-18 would also enable us to regulate the polarization of human M2 pro-tumoral macrophages towards M1 antitumoral macrophages (**Fig. S6a** and following the gating strategy described in **Fig. S6b**). We therefore generated iTRUCK19.18 cells and isolated the monocytes from the same donor, which were then differentiated into an M2 phenotype (**Fig. 6A**). Following co-culture of iTRUCK19.18 and M2 macrophages (CD206+CD11c-), we observed the polarization of the macrophages towards M1 (CD206-CD11c+), antitumoral phenotype only in the presence of Dox (**Fig. 6b**).

**Fig. 6.**
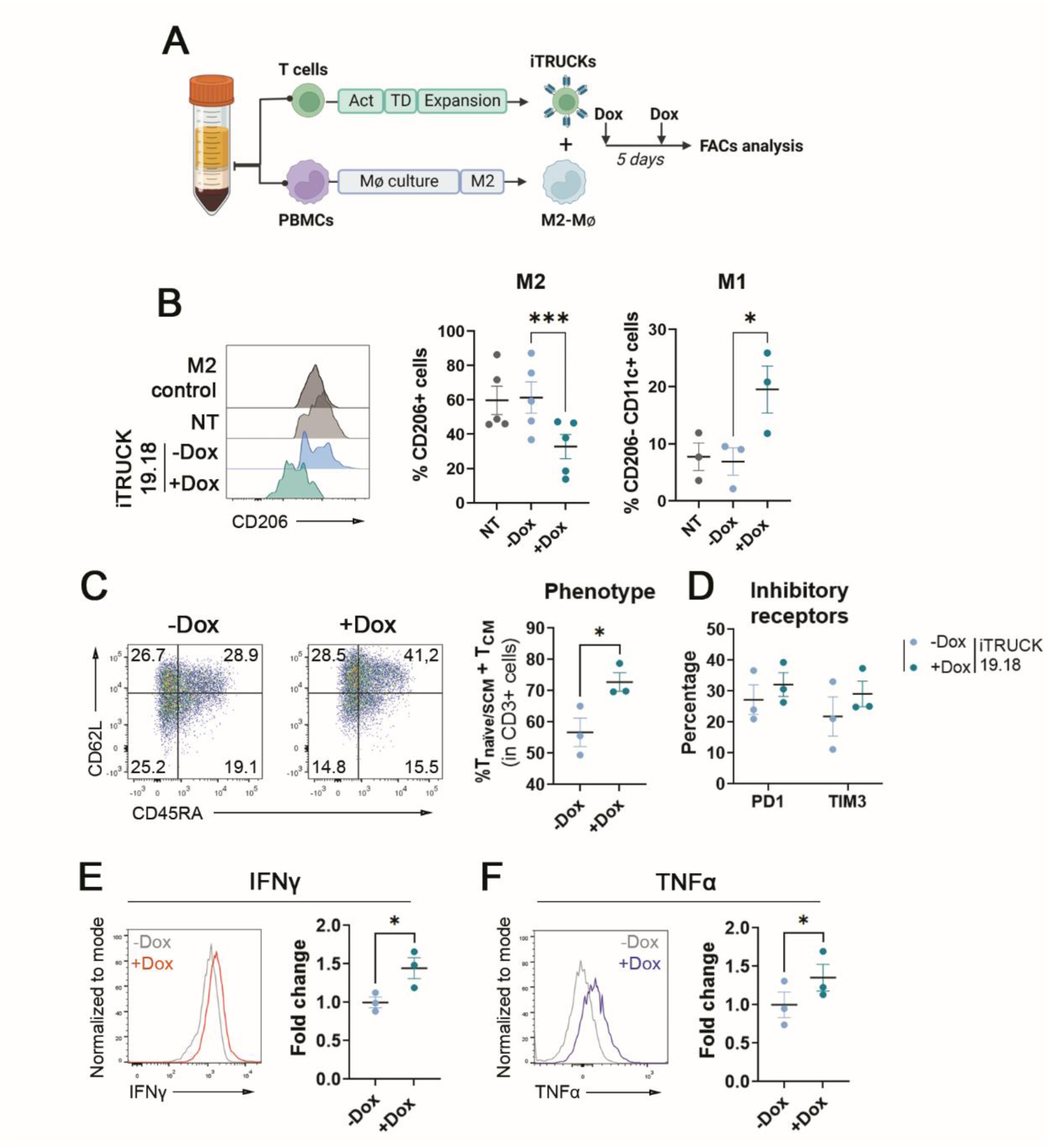
Dox addition to iTRUCK19.18 cells induces polarization of primary human M2 macrophages towards an M1 phenotype. **a** Experimental diagram of the generation of iTRUCK19.18 and M2-polarized macrophages from the same donor. Briefly, non-adherent cells from the PBMCs layer were activated (Act) with αCD3/C28, transduced (TD) and expanded whereas adherent cells from the mononuclear phase was cultured and polarized as described in M&M. Then, NT and iTRUCK19.18 (previously cultured in the absence of Dox) were co-cultured the M2-macrophages during 5 days until FACS analysis. Dox (50 ng/ml) was added at the beginning of the co-culture. **b** Left: representative histograms corresponding to CD206 expression of macrophages cocultured with the different groups of T cells. Center and right: Percentage of CD206+ (center) and M1 (right) macrophages when co-cultured with NT cells, iTRUCK19.18 cells without and with Dox, and with MIA PaCa2 CD19+ cells at ratio 1:1 MiaPaca vs CAR-T (M2: n=5; M1: n=3). **c** Representative dot plots and quantification of the proportion of T_naïve/SCM_ and T_CM_ cells of iTRUCK19.18 cells in the presence of macrophages with and without Dox after 5 days of the co-culture (n=3). **d** Percentage of PD1 and TIM3 positive cells of iTRUCK19.18 cells after the co-culture in the presence (dark blue) or absence (light blue) of Dox (n=3). **e** Representative histograms and fold change of IFNγ expression (compared to those without Dox) by iTRUCK19.18 cells in co-culture with macrophages with and without Dox (n=3). **f** Representative histograms and fold change of TNFα expression by iTRUCK19.18 cells in co-culture with macrophages with and without Dox (n=3). *p<0.05, ***p<0.001 (two-tailed paired t test for B; one-tailed paired t test for C, D, E and F).

Furthermore, we evaluated how the release of IL-18 affected the state of iTRUCK19.18 cells when co-cultured with macrophages. Interestingly, we observed again that iTRUCK19.18 cells exposed to Dox displayed a less differentiated phenotype, with a higher proportion of T_naïve/SCM_ and T_CM_ cells, in contrast to cells without Dox (**Fig. 6c**). No significant differences were found in the expression of PD1 and TIM3 (**Fig. 6d**). Of note, we validated that the induction of IL-18 in this microenvironment led to a Dox-dependent increase in the production of IFNγ (**Fig. 6e**) and TNFα (**Fig. 6f**) by iTRUCK19.18 cells.

All together this data indicates that iTRUCK19.18 cells can be externally controlled not only to enhance their potency but also to act on other immune cells, such as macrophages, to potentially re-shape the TME.

## Discussion

Clinical experience has highlighted that CAR-T therapy still has significant room for improvement when applied to patients with aggressive/resistant neoplasms. Conventional CAR-T cells have not demonstrated sufficient efficacy in many cancers, leading to frequent relapses in liquid neoplasms and an almost complete lack of therapeutic effect in solid tumors. One of the most promising approaches to enhance the effectiveness of CAR-T therapy in recurrent neoplasms is the development of 4^th^ generation CAR-T cells (TRUCKs). In clinical trials, these TRUCKs have shown promising results by overexpressing cytokines such as IL-12 [NCT02498912], IL-15 [NCT03579888] and IL-18 [NCT04684563], which exert a strong immune-modulating action and allow for the restructuring of the TME. TME plays a pivotal role in causing premature dysfunction of CAR-T cells. Consequently, strategies aimed at mitigating its immunosuppressive actions are emerging as interesting alternatives to improve treatment outcomes.

Despite the promising therapeutic effect of TRUCKs that overexpress cytokines in hard-to-treat tumors, their continuous immunomodulatory action can also lead to serious toxicities. Sustained administration of IL-12 and IL-15 has been linked to severe toxicity events (38, 39), while the constitutive expression of IL-18 has been associated not only with specific toxicity events (30) but also with the onset of autoimmune disorders (31) and IFNγ-independent toxicities (32). Therefore, it is essential to control the expression of these cytokines in CAR-T cells to achieve an appropriate balance between efficacy and safety.

The primary mechanism for controlling these cytokines is the use of activation-inducible promoters, particularly the NFAT promoter, which has been applied to regulate IL-12 or IL-18 [NCT03542799] (26) among others. While yielding promising results, there have been reported cases where the NFAT promoter has failed to effectively regulate IL-12 expression, resulting in *in vivo* toxicity (10, 11). This is because this promoter is activated in response to T cell activation, which may not be exclusively dependent on the antigenic recognition by the CAR. As an alternative, approaches using Tet-On systems, which enable inducible gene expression through doxycycline, have shown promising results, although their use have been relatively underexplored. Traditional Tet-On systems present certain significant limitations. First, the most commonly used doxycycline induction systems (Tet-On-3G system by Takara Bio, or the system proposed by Alsaieedi and colleagues) require very high concentrations of doxycycline (in this latter case, 2 mg/ml) for *in vivo* induction (11). This could potentially contribute to the development of long-term bacterial resistance due to prolonged or intermittent exposure (15, 40). Additionally, most of these systems require transactivator proteins (rtTA). These bacterial/viral chimeras are necessary to trigger transgene transcription. This point is crucial because important safety considerations must be made since these elements appear to hijack transcription factors in a non-expected manner. When combined with their ability to bind to *pseudo*-tetO sites throughout the genome, they can cause undesired and nonspecific transactivation of genes (16–18, 41). In addition, Schmitt *et al*. have described alterations due to transactivators in activated, memory and regulatory splenic T cells subsets in transgenic mice carrying a Tet-On transactivator after only 6 days in the presence of Dox (42). Altogether, the use of transactivators seems to strongly hinder safe clinical applications. To prevent these potential toxicities for safer ATPMs, we used the Lent-On-Plus (LOP) system to generate αCD19 CAR-T cells able to induce IL-18 expression under doxycycline. This system, which does not rely on transactivator proteins, allows for a closer approach to clinical application. Moreover, the system allows *in vivo* induction under ultra-low doses of doxycycline at nanogram level (23). To our knowledge, we present the first CAR-T cells able to exogenously induce IL-18 expression, and the first doxycycline-inducible/transactivator-free TRUCK (iTRUCK19.18).

iTRUCK19.18 cells generation through co-transduction with CAR19 and LOP18 LVs leads to heterogeneous cell populations. Although obtaining more homogeneous populations would be ideal (43), in the case of IL-18, its importance is diminished since CAR+IL18-cells exhibit antitumor activity *per se*, while CAR-IL18+ cells provide support to all T cells, whether they carry the CAR or not, as all T cells express IL-18 receptors. This bystander effect enhances their antitumor activity, as previously demonstrated by Hu and colleagues (28). Moreover, using two independent vectors provides versatility to the system, enabling its application in combination with any synthetic CAR, patient-derived TCR or even on TILs without the need to modify the lentiviral backbone. It is important to note that, despite the absence of clear evidence for a superior method of expressing multiple transgenes on T cells, co-transduction has been demonstrated as safe (44), and has produced solid results, even better than using bicistronic vectors in some cases (45, 46), recently reaching the clinical stage (47) (NCT02443831). In the same direction, we observed that the production of iTRUCK19.18 cells through co-transduction does not significantly affect parameters related to activation and phenotype compared to the production of conventional CAR-T cells. Co-transduction overcomes the technical difficulties of using bicistronic vectors that have led to poor clinical data, and allows lower costs compared to two full cell products manufacturers for pooled co-infusion.

It is interesting to note that the secretion of IL-18 by iTRUCK19.18 cells is detected when both the cells are activated and doxycycline is added, following the natural process of IL-18 processing and secretion observed in other cell types (36). Interestingly, in the absence of activation but in the presence of doxycycline, intracellular pro-IL-18 was detected, and functional effects on T cells were observed. These results suggest that, even without activation, iTRUCK19.18 cells can express low levels of IL-18 upon Dox addition, inducing certain functional effects. Importantly, these findings strongly indicate that the IL-18 produced by iTRUCK19.18 cells is bioactive, and that the combination of the double safety mechanism activation/Dox allows for the generation of a safer system compared with the standardized use of NFAT-based promoters.

In order to target different aggressive tumor contexts, we employed iTRUCK19.18 cells in both liquid and CD19-engineered solid tumor models, as well as in patient tumor samples. Analysis in lymphoma model (Namalwa) demonstrated that iTRUCK19.18 cells exhibit superior antitumoral activity compared to conventional CAR-T cells (in line with the observations of Hu *et al.* and Avanzi *et al.* targeting CD19+ neoplasms) (28, 29), and this activity can be controlled using Dox. After confirming the effectiveness of our system in an aggressive liquid neoplasm model, we assessed its activity in a solid tumor model to verify that the effectiveness is not dependent on the tumor type. Once again, we observed a significant increase in antitumoral potency using a CD19+ pancreatic tumor model (in line with studies reporting increased antitumoral activity of IL-18 releasing CAR-T cells against solid tumors) (26, 48). Consistent with the lymphoma model, we found a less differentiated phenotype, leading to heightened antitumoral efficacy. Apart from the augmented activation, we hypothesized that this improved phenotype and retained cellular exhaustion contribute to enhanced T cell fitness, ultimately resulting in superior antitumoral potency, aligning with recent findings reported by Jaspers *et al* (48). Additional studies are warranted to elucidate the inherent mechanistic factors linked to the exogenous expression of IL-18 on T cells. This phenomenon likely correlates with a decrease in markers indicative of terminal differentiation such as TOX.

The generation of iTRUCK19.18 cells from patient T cells confirmed the feasibility of co-transduction to achieve sufficient expression levels of both transgenes without any significant alteration on T cell fitness and provides a clinically closer approach of the IL-18-derived increased antitumor potency of iTRUCK19.18.

Our findings clearly showed the positive effects of Dox-induced IL-18 on T cells, resulting in an increased antitumoral activity of iTRUCK19.18 cells. However, IL-18 has a plethora of biological actions on different cells that are crucial over the TME. We therefore used an *in vitro* model for the same donor to demonstrate that pro-tumoral human M2 tumor-associated macrophages can be polarized to an antitumoral M1 phenotype by iTRUCK19.18 cells in a Dox-dependent manner. This is consistent with what was observed by Chmielewski and colleagues in a murine model (26). Overall, this suggests that IL-18 may potentially reshape the TME, enhancing the immune-activating and cytotoxic function of CAR-T cells.

Our results indicate that controlling the release of IL-18 by CAR-T cells also allows for controlling their antitumor potency in different tumor contexts. Cytokine control through an ultra-low dose Dox-inducible system free of transactivators (LOP) allows for the generation of a safe and more effective TRUCKs product, representing an alternative to conventional CAR-T cells for treating patients with aggressive type B neoplasms that require increased potency without compromising safety.

## Conclusions

In summary, we have demonstrated that the LOP platform allows the development of 4th generation, inducible CAR-T cells (iTRUCKS) in the absence of transactivators, that can be controlled under ultra-low doses of doxycycline. Using this platform, we have generated iTRUCK19.18 from healthy donors and B-cell neoplasms patients and demonstrated it superior activity on a lymphoma model, in patient’s tumor cells, and in a CD19+ solid tumor model. Moreover, we demonstrated that iTRUCK19-18 exhibits dual safety control mechanisms, necessitating both T cell activation and the presence of doxycycline for IL-18 secretion. This dual control not only enhances T cell potency but also induces the polarization of pro-tumoral M2 macrophages towards an antitumoral phenotype (M1). Based on these results, we suggest iTRUCK19.18 as a novel alternative for treating type-B lymphomas that are resistant to standard CAR-T cells. Additionally, we anticipate the prospective utilization of LOP18 as an inducible booster to augment the potency of other CAR-T cells in a regulated manner. Finally, we advocate for the incorporation of the LOP platform as an additional tool in the arsenal for developing more potent and controllable advanced therapy medicinal products (iATMPs).

## Availability of data and materials

All data are available in the main text or in the supplemental information. LOP-18 LV is available under a material transfer agreement with LentiStem Biotech.

## Acknowledgments

We acknowledge Dr. Manel Juan and Dra. Maria Castella for kindly providing CAR19 LV. We also thanks Dra. Paulina Rybakowska, Dra. Araceli Aguilar and Paula Heredia for their support with human samples; Ana Fernández-Ibáñez for her support with the IVIS Spectrum Analyzer and all supporting Units from GENYO.

## Funding

Instituto de Salud Carlos III (ISCIII) and the European Regional Development Fund (FEDER): Research grants, **PI21/00298.** (FM)

Instituto de Salud Carlos III (ISCIII) – NextGenerationEU funds - actions of the Recovery and Resilience Mechanism: Red TerAv **RD21/ 0017/0004.** (FM, JAM)

Ministerio de Ciencia e innovación (MICIN). Plan de Recuperación, transformación y resilencia, Centro para el Desarrollo Tecnológico Industrial (CDTI) and European Union-Next Generation EU: Research grants **00123009/SNEO-20191072 (FM)**, **PMPTA22/00060 (FM)** and **DIN2018-010180.** (PJL)

Consejería de Salud y Familias (CSyF) -Junta de Andalucía - FEDER/European Cohesion Fund (FSE) for Andalucía: Grants: 2**016000073332-TRA**, **CARTPI-0001-201**, **PECART-0031-2020, PECART-0027-2020** y **CAR-T 2019 00400200101918 (Red RANTECAR**). (FM, JAM)

Consejería de Economía, conocimiento, empresa y Universidad. Grant **A-CTS-235-UGR18**. (FM)

Ministerio de Ciencia e innovación (MICIN) – líneas estratégicas: Grant **PLEC2021-008094** (FM, JAM). Fellowship Garantía Juvenil (PEJ2018-001760-A) (MCG).

Chair “Doctors Galera-Requena in cancer stem cell research” (**CMC-CTS963**) (JAM).

## Author contributions

Conceptualization: FM, MTM, PJL, NMP

Methodology: PJL, MTM, NMP, FM, JAM

Investigation: PJL, MTM, NMP, CBJ, MCG, KP, PM, AHB, CGL, SANM

Resources: FM, JMB, PAGS, CH

Writing - Original Draft: PJL, MTM, FM

Writing - Review & Editing: FM, MTM, PJL, FJME

Supervision: MTM, FM

Funding acquisition: FM, JAM

## Competing interests

FM, PM are inventors of the patents entitled “Lent-on-plus system for conditional expression in human stem cells” (PCT/EP2017/078246) and “Insulator to improve gene transfer vectors” (PCT/EP2014/055027). FM, MTM and JAM are partners of LentiStem Biotech. PJL, MTM, and CBJ are contractually associated with LentiStem Biotech, a spin-off company that holds the license of the above-mentioned patents. All other authors declare they have no competing interests.

## List of abbreviations

AICD: Activation-induced cell death
ATMPs: Advanced Therapy Medicinal Products
CAR: Chimeric Antigen Receptor
CLL: Chronic Lymphocytic Leukemia
Dox: Doxycycline
FasL: Fas Ligand
GPCR: G Protein-Coupled Receptor
GFP-Nluc: Green Fluorescent Protein-Nanoluciferase
GVHD: Graft-versus-host-disease
IFNγ: Interferon gamma
IL-18: Interleukin 18
LOP: Lent-On-Plus
LV: Lentiviral Vector
MOI: Multiplicity of Infection
MZL: Marginal Zone Lymphoma
NK: Natural Killer
NFAT: Nuclear Factor of Activated T cell
NSG: NOD/scid-IL-2Rnull mice
NT: Non-transduced T cells
PDAC: Pancreatic Ductal Adenocarcinoma
P/S: Penicillin/Streptomycin
R/R: Relapsed/Refractory
TAMs: Tumor Associated Macrophages
Tet-On: Tetracycline Operon On
TetR: Tetracycline Repressor
TME: Tumor Microenvironment
TNFα: Tumor Necrosis Factor Alpha
TRAIL: TNF-related apoptosis-inducing ligand
TRUCKs: T cells Redirected for Antigen-Unrestricted Cytokine-initiated Killing
rtTA: Transactivator protein

## Notes

### Competing Interest Statement

FM and PM are inventors of the patents entitled 'Lent-on-plus system for conditional expression in human stem cells' (PCT/EP2017/078246) and 'Insulator to improve gene transfer vectors' (PCT/EP2014/055027). FM, MTM and JAM are partners of LentiStem Biotech. PJL, MTM, and CBJ are contractually associated with LentiStem Biotech, a spin-off company that holds the license of the above-mentioned patents. All other authors declare they have no competing interests.

## References

1. Melenhorst JJ, Chen GM, Wang M, Porter DL, Chen C, Collins MA, et al. Decade-long leukaemia remissions with persistence of CD4+ CAR T cells. Nature. 2022;602(7897):503–9.

2. Gu T, Zhu M, Huang H, Hu Y. Relapse after CAR-T cell therapy in B-cell malignancies: challenges and future approaches. J Zhejiang Univ Sci B. 2022;23(10):793–811.

3. Todorovic Z, Todorovic D, Markovic V, Ladjevac N, Zdravkovic N, Djurdjevic P, et al. CAR T Cell Therapy for Chronic Lymphocytic Leukemia: Successes and Shortcomings. Curr Oncol. 2022;29(5):3647–57.

4. Safarzadeh Kozani P, Safarzadeh Kozani P, Ahmadi Najafabadi M, Yousefi F, Mirarefin SMJ, Rahbarizadeh F. Recent Advances in Solid Tumor CAR-T Cell Therapy: Driving Tumor Cells From Hero to Zero? Front Immunol. 2022;13:795164.

5. Mazinani M, Rahbarizadeh F. CAR-T cell potency: from structural elements to vector backbone components. Biomark Res. 2022;10(1):70.

6. Fraietta JA, Lacey SF, Orlando EJ, Pruteanu-Malinici I, Gohil M, Lundh S, et al. Determinants of response and resistance to CD19 chimeric antigen receptor (CAR) T cell therapy of chronic lymphocytic leukemia. Nat Med. 2018;24(5):563–71.

7. Li J, Li W, Huang K, Zhang Y, Kupfer G, Zhao Q. Chimeric antigen receptor T cell (CAR-T) immunotherapy for solid tumors: lessons learned and strategies for moving forward. J Hematol Oncol. 2018;11(1):22.

8. Chmielewski M, Abken H. TRUCKS, the fourth-generation CAR T cells: Current developments and clinical translation. ADVANCES IN CELL AND GENE THERAPY. 2020;3(3):e84.

9. Chmielewski M, Abken H. TRUCKs: the fourth generation of CARs. Expert Opin Biol Ther. 2015;15(8):1145–54.

10. Zhang L, Morgan RA, Beane JD, Zheng Z, Dudley ME, Kassim SH, et al. Tumor-infiltrating lymphocytes genetically engineered with an inducible gene encoding interleukin-12 for the immunotherapy of metastatic melanoma. Clin Cancer Res. 2015;21(10):2278–88.

11. Alsaieedi A, Holler A, Velica P, Bendle G, Stauss HJ. Safety and efficacy of Tet-regulated IL-12 expression in cancer-specific T cells. Oncoimmunology. 2019;8(3):1542917.

12. Wang D. The essential role of G protein-coupled receptor (GPCR) signaling in regulating T cell immunity. Immunopharmacol Immunotoxicol. 2018;40(3):187–92.

13. Condotta SA, Richer MJ. The immune battlefield: The impact of inflammatory cytokines on CD8+ T-cell immunity. PLoS Pathog. 2017;13(10):e1006618.

14. Agnellini P, Wolint P, Rehr M, Cahenzli J, Karrer U, Oxenius A. Impaired NFAT nuclear translocation results in split exhaustion of virus-specific CD8+ T cell functions during chronic viral infection. Proc Natl Acad Sci U S A. 2007;104(11):4565–70.

15. Tristan-Manzano M, Justicia-Lirio P, Maldonado-Perez N, Cortijo-Gutierrez M, Benabdellah K, Martin F. Externally-Controlled Systems for Immunotherapy: From Bench to Bedside. Front Immunol. 2020;11:2044.

16. Morimoto M, Kopan R. rtTA toxicity limits the usefulness of the SP-C-rtTA transgenic mouse. Dev Biol. 2009;325(1):171–8.

17. Perl AK, Zhang L, Whitsett JA. Conditional expression of genes in the respiratory epithelium in transgenic mice: cautionary notes and toward building a better mouse trap. Am J Respir Cell Mol Biol. 2009;40(1):1–3.

18. Whitsett JA, Perl AK. Conditional control of gene expression in the respiratory epithelium: A cautionary note. Am J Respir Cell Mol Biol. 2006;34(5):519–20.

19. Sisson TH, Hansen JM, Shah M, Hanson KE, Du M, Ling T, et al. Expression of the reverse tetracycline-transactivator gene causes emphysema-like changes in mice. Am J Respir Cell Mol Biol. 2006;34(5):552–60.

20. Benabdellah K, Gutierrez-Guerrero A, Cobo M, Munoz P, Martin F. A chimeric HS4-SAR insulator (IS2) that prevents silencing and enhances expression of lentiviral vectors in pluripotent stem cells. PLoS One. 2014;9(1):e84268.

21. Benabdellah K, Cobo M, Munoz P, Toscano MG, Martin F. Development of an all-in-one lentiviral vector system based on the original TetR for the easy generation of Tet-ON cell lines. PLoS One. 2011;6(8):e23734.

22. Benabdellah K, Muñoz P, Cobo M, Gutierrez-Guerrero A, Sánchez-Hernández S, Garcia-Perez A, et al. Lent-On-Plus Lentiviral vectors for conditional expression in human stem cells. Scientific Reports. 2016;6(1):37289.

23. Tristán-Manzano M, Maldonado-Pérez N, Justicia-Lirio P, Cortijo-Gutierréz M, Tristán-Ramos P, Blanco-Benítez C, et al. Lentiviral vectors for inducible, transactivator-free advanced therapy medicinal products: Application to CAR-T cells. Molecular Therapy - Nucleic Acids. 2023;32:322–39.

24. Hoshino T, Wiltrout RH, Young HA. IL-18 is a potent coinducer of IL-13 in NK and T cells: a new potential role for IL-18 in modulating the immune response. J Immunol. 1999;162(9):5070–7.

25. Ohtsuki T, Micallef MJ, Kohno K, Tanimoto T, Ikeda M, Kurimoto M. Interleukin 18 enhances Fas ligand expression and induces apoptosis in Fas-expressing human myelomonocytic KG-1 cells. Anticancer Res. 1997;17(5A):3253–8.

26. Chmielewski M, Abken H. CAR T Cells Releasing IL-18 Convert to T-Bet(high) FoxO1(low) Effectors that Exhibit Augmented Activity against Advanced Solid Tumors. Cell Rep. 2017;21(11):3205–19.

27. Olivera I, Bolanos E, Gonzalez-Gomariz J, Hervas-Stubbs S, Marino KV, Luri-Rey C, et al. mRNAs encoding IL-12 and a decoy-resistant variant of IL-18 synergize to engineer T cells for efficacious intratumoral adoptive immunotherapy. Cell Rep Med. 2023;4(3):100978.

28. Hu B, Ren J, Luo Y, Keith B, Young RM, Scholler J, et al. Augmentation of Antitumor Immunity by Human and Mouse CAR T Cells Secreting IL-18. Cell Rep. 2017;20(13):3025–33.

29. Avanzi MP, Yeku O, Li X, Wijewarnasuriya DP, van Leeuwen DG, Cheung K, et al. Engineered Tumor-Targeted T Cells Mediate Enhanced Anti-Tumor Efficacy Both Directly and through Activation of the Endogenous Immune System. Cell Rep. 2018;23(7):2130–41.

30. Breman E, Walravens A-S, Gennart I, Velghe A, Nguyen T, Violle B, et al. 107 Armoring NKG2D CAR T cells with IL-18 improves in vivo anti-tumor activity. Journal for ImmunoTherapy of Cancer. 2021;9(Suppl 2):A118-A.

31. Tsutsumi N, Yokota A, Kimura T, Kato Z, Fukao T, Shirakawa M, et al. An innate interaction between IL-18 and the propeptide that inactivates its precursor form. Sci Rep. 2019;9(1):6160.

32. Nakamura S, Otani T, Ijiri Y, Motoda R, Kurimoto M, Orita K. IFN-γ-Dependent and -Independent Mechanisms in Adverse Effects Caused by Concomitant Administration of IL-18 and IL-12. The Journal of Immunology. 2000;164(6):3330–6.

33. Frecha C, Toscano MG, Costa C, Saez-Lara MJ, Cosset FL, Verhoeyen E, et al. Improved lentiviral vectors for Wiskott-Aldrich syndrome gene therapy mimic endogenous expression profiles throughout haematopoiesis. Gene Ther. 2008;15(12):930–41.

34. Larson RC, Kann MC, Bailey SR, Haradhvala NJ, Llopis PM, Bouffard AA, et al. CAR T cell killing requires the IFNgammaR pathway in solid but not liquid tumours. Nature. 2022;604(7906):563–70.

35. Castella M, Boronat A, Martin-Ibanez R, Rodriguez V, Sune G, Caballero M, et al. Development of a Novel Anti-CD19 Chimeric Antigen Receptor: A Paradigm for an Affordable CAR T Cell Production at Academic Institutions. Mol Ther Methods Clin Dev. 2019;12:134–44.

36. van de Veerdonk FL, Netea MG, Dinarello CA, Joosten LA. Inflammasome activation and IL-1beta and IL-18 processing during infection. Trends Immunol. 2011;32(3):110–6.

37. Tristán-Manzano M, Maldonado-Pérez N, Justicia-Lirio P, Muñoz P, Cortijo-Gutiérrez M, Pavlovic K, et al. Physiological lentiviral vectors for the generation of improved CAR-T cells. Molecular Therapy - Oncolytics. 2022;25:335–49.

38. Leonard JP, Sherman ML, Fisher GL, Buchanan LJ, Larsen G, Atkins MB, et al. Effects of single-dose interleukin-12 exposure on interleukin-12-associated toxicity and interferon-gamma production. Blood. 1997;90(7):2541–8.

39. Miller JS, Morishima C, McNeel DG, Patel MR, Kohrt HEK, Thompson JA, et al. A First-in-Human Phase I Study of Subcutaneous Outpatient Recombinant Human IL15 (rhIL15) in Adults with Advanced Solid Tumors. Clin Cancer Res. 2018;24(7):1525–35.

40. Grossman TH. Tetracycline Antibiotics and Resistance. Cold Spring Harb Perspect Med. 2016;6(4):a025387.

41. Hackl H, Rommer A, Konrad TA, Nassimbeni C, Wieser R. Tetracycline regulator expression alters the transcriptional program of mammalian cells. PLoS One. 2010;5(9):e13013.

42. Schmitt A, Schulze-Osthoff K, Hailfinger S. Correspondence: T cells are compromised in tetracycline transactivator transgenic mice. Cell Death Differ. 2018;25(3):634–6.

43. Cordoba S, Onuoha S, Thomas S, Pignataro DS, Hough R, Ghorashian S, et al. CAR T cells with dual targeting of CD19 and CD22 in pediatric and young adult patients with relapsed or refractory B cell acute lymphoblastic leukemia: a phase 1 trial. Nat Med. 2021;27(10):1797–805.

44. Frimpong K, Spector SA. Cotransduction of nondividing cells using lentiviral vectors. Gene Ther. 2000;7(18):1562–9.

45. Spiegel JY, Patel S, Muffly L, Hossain NM, Oak J, Baird JH, et al. CAR T cells with dual targeting of CD19 and CD22 in adult patients with recurrent or refractory B cell malignancies: a phase 1 trial. Nature Medicine. 2021;27(8):1419–31.

46. Bachiller M, Dobaño-López C, Rodríguez-García A, Castellsagué J, Gimenez-Alejandre M, Antoñana-Vildosola A, et al. Co-Transduced CD19/BCMA Dual-Targeting CAR-T Cells for the Treatment of Non-Hodgkin Lymphoma. Blood. 2022;140(Supplement 1):7386–7.

47. Ghorashian S, Lucchini G, Richardson R, Nguyen K, Terris C, Guvenel A, et al. CD19/CD22 targeting with co-transduced CAR T-cells to prevent antigen negative relapse after CAR T-cell therapy of B-ALL. Blood. 2023.

48. Jaspers JE, Khan JF, Godfrey WD, Lopez AV, Ciampricotti M, Rudin CM, et al. IL-18-secreting CAR T cells targeting DLL3 are highly effective in small cell lung cancer models. J Clin Invest. 2023.

