## Supplementary materials for "*First-in-class* Transactivator-Free, Doxycycline-inducible IL-18-engineered CAR-T cells for relapsed/refractory B-cell lymphomas"

[†]Share first authorship

[‡]Share co-senior authorship

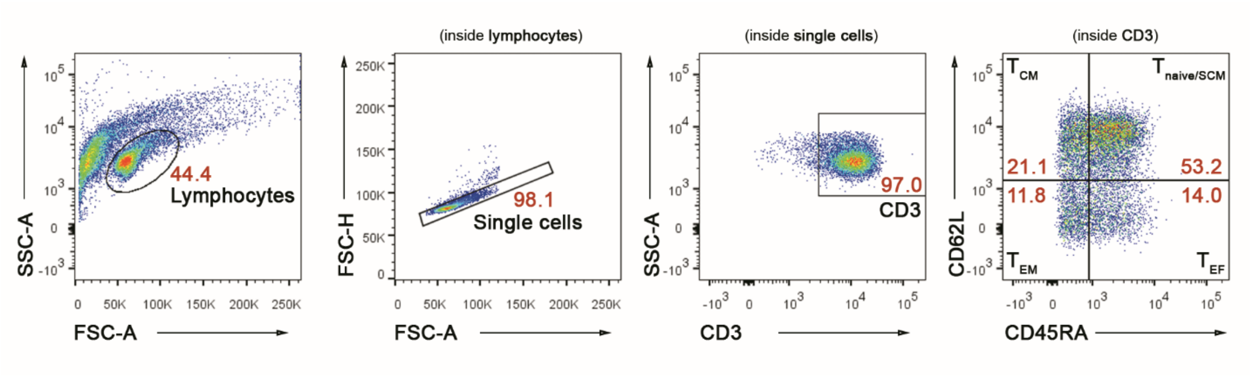


**Fig. S1** Gating strategy for the analysis of T cell phenotype. After selecting the lymphocytes gate regarding to FSC and SSC, doubles exclusion and confirming CD3, four populations were established according to the expression of CD45RA and CD62L. $T_{naive/SCM}$: CD45RA+CD62L+; $T_{CM}$: CD45RA-CD62L-; $T_{EM}$: CD45RA-CD62L-; $T_{EF}$: CD45RA+CD62L-.

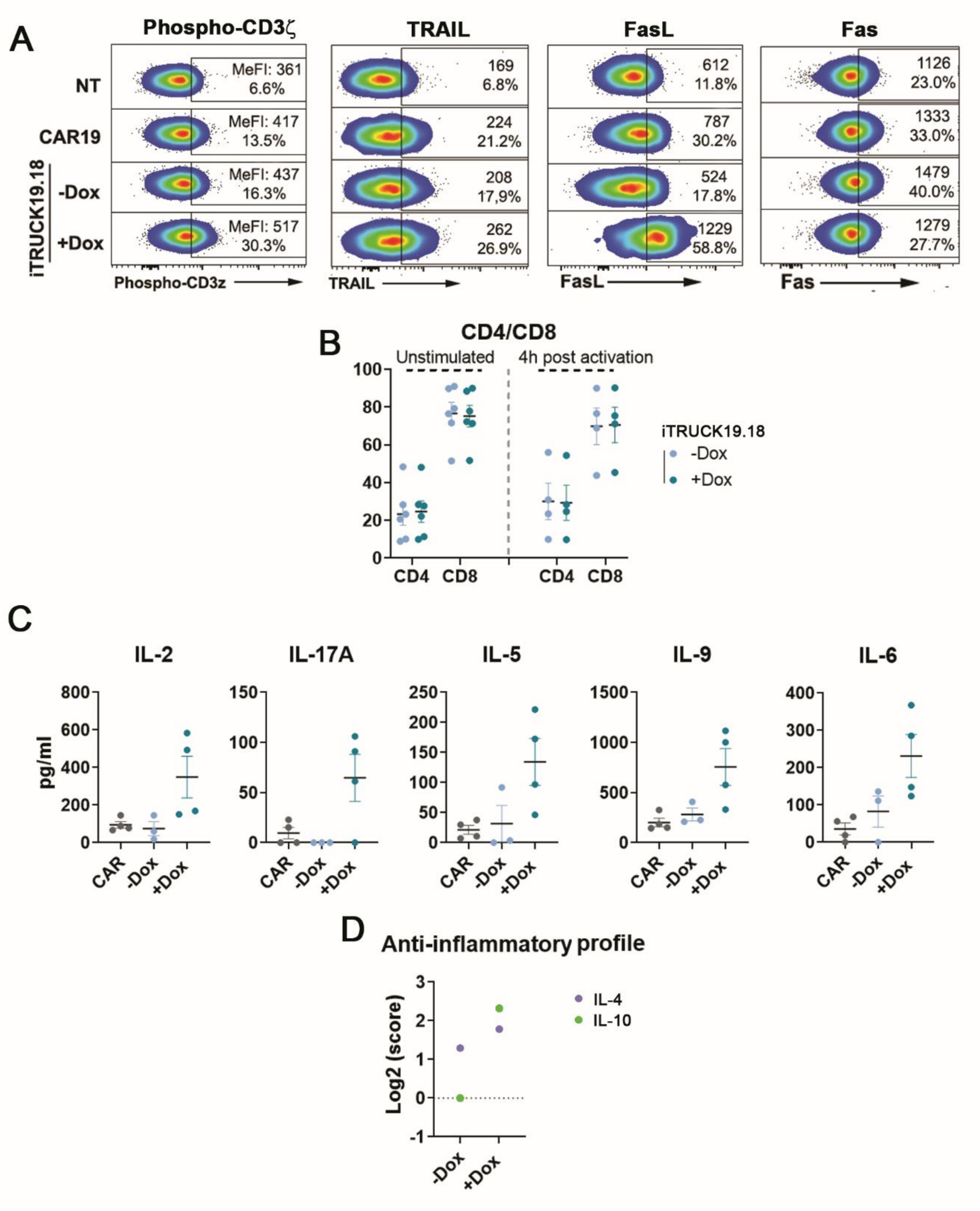


**Fig. S2** Characterization of iTRUCK19.18 based on AICD markers and their production of immunomodulatory cytokines at basal state. **a** Representative dot-plots of phosphorylated-CD3ζ, TRAIL, FasL and Fas of NT, CAR19, iTRUCK19.18 cells (-Dox and +50 ng/ml Dox) after manufacturing at basal conditions. **b** Percentage of CD4/CD8 at basal state (n=6) (left) and 4h post stimulation via CD3/CD28 (n=4) (right) of iTRUCK19.18 with (50 ng/ml) and without Dox. **c** Quantification of proinflammatory cytokine secretion by iTRUCK19.18 cells (from left to right:

IL-2, IL-17A, IL-5, IL-9, and IL-6) (n=2). **d** Anti-inflammatory index of by iTRUCK19.18 cells (IL-4 and IL-10) (n=2) (Dox: 50 ng/ml).

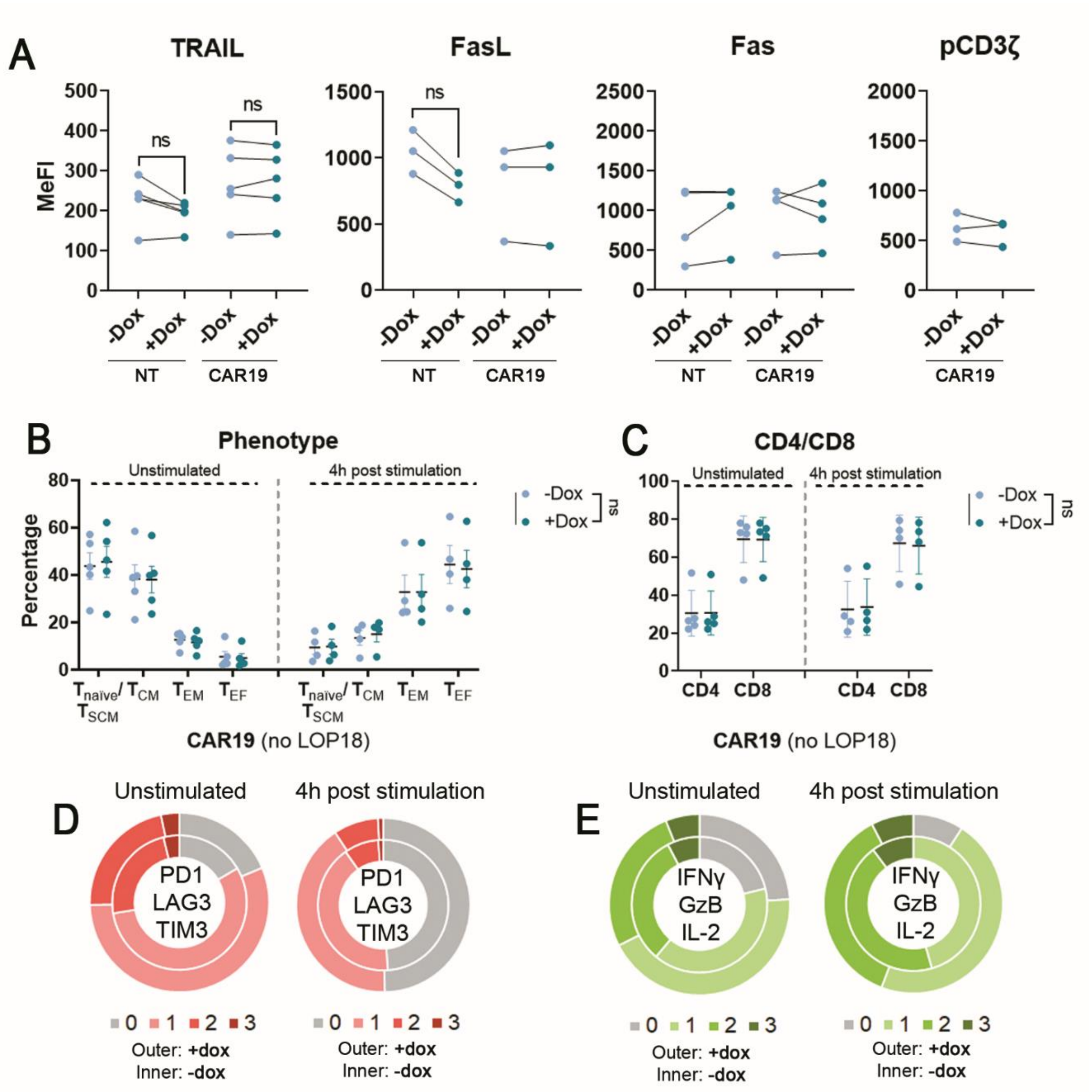


**Fig. S3** Doxycycline does not alter the expression of the main markers used to characterize iTRUCK19.18 cells. **a** Expression of AICD markers and phosphorylated-CD3ζ (from left to right) in NT and CAR19 cells at basal state in the presence (50 ng/ml) or absence of Dox at basal conditions after manufacturing. TRAIL (n=5), FasL (n=3), Fas (n=4) and pCD3ζ (n=3) (two-tailed paired t test). **b** Percentage of positive cells of $T_{\text{Naïve/SCM}}$, $T_{CM}$, $T_{EM}$ and $T_{EF}$ at basal state (n=5) (left) and 4h post stimulation (n=4) (right) of CAR19 cells with (50 ng/ml) and without Dox. **c** Percentage of CD4 and CD8 of CAR19 cells with (50 ng/ml) and without Dox at basal state (n=5) (left) and 4h post stimulation (n=4) (right). **d** Pie charts showing the proportion of CAR19 cells with (50 ng/ml) (outer circles) and without (inner circles) Dox expressing 0, 1, 2, or 3 inhibitory receptors (PD1, LAG3, and TIM3) at basal state (n=4) (left) and 4h after stimulation (n=4) (right). **e** Pie charts showing the proportion of CAR19 cells with (50 ng/ml) (outer circles) and without (inner circles) Dox expressing 0, 1, 2, or 3 activation markers (IFNγ, Granzyme B, and IL-2) at basal state (n=4) (left) and 4h post stimulation (n=4) (right). Activation was performed using T cell TransAct (Miltenyi) (via CD3/CD28). ns: non-significant (two-tailed paired t test for A, one-tailed paired t test for B and C).

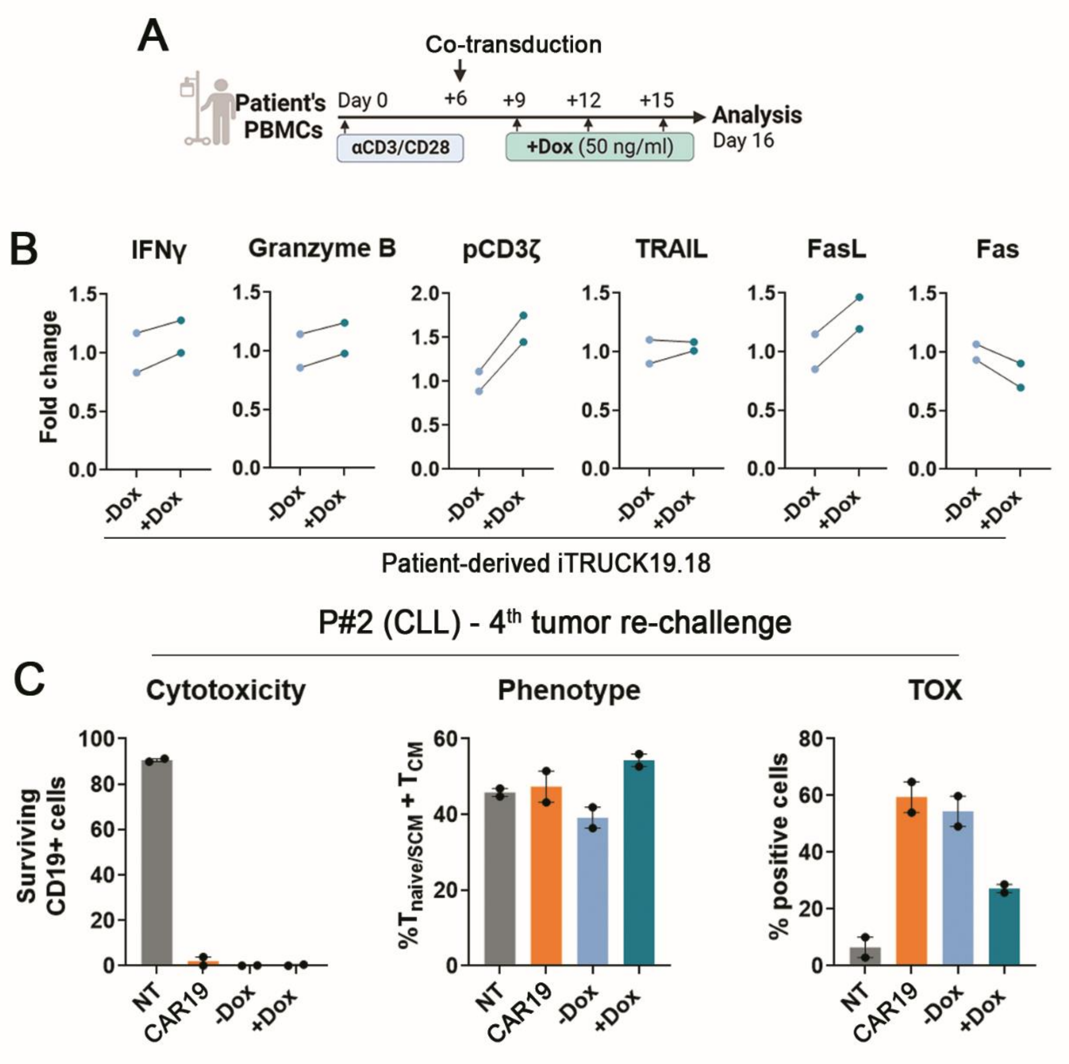


**Fig. S4** Generation and characterization of patient-derived iTRUK19.18: cytotoxicity, phenotype and TOX expression of patient 2-derived iTRUCK19.18 cells after 5 tumor encounters with their own tumor. **a** Experimental diagram of iTRUCK19.18 generation from patient's PBMCs. Briefly, after 6 days in the presence of αCD3/CD28, enriched T cells were transduced with CAR19 and LOP18 LVs as described in M&M. **b** Fold-change (relative to -Dox) of activation-related markers expression of patient-derived iTRUCK19.18 cells without and with (50 ng/ml) Dox treatment under basal conditions (from left to right: IFNγ, Granzyme B, phospho-CD3ζ, TRAIL, FasL, and Fas) (n=2, P#2 and P#3). **c** Graph depicting surviving tumor cells (left), proportion of $T_{naive/SCM}$ and $T_{CM}$ cells (center), and TOX expression (right), all after the 4$^{th}$ tumor re-challenge (n=1, P#2) (Dox=50 ng/ml).

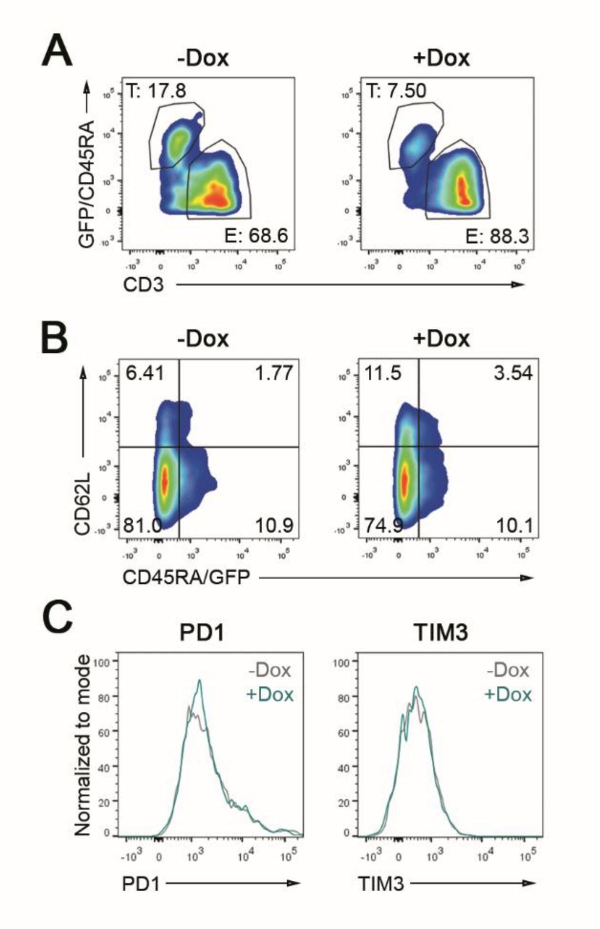


**Fig. S5** Representative dot-plots of iTRUCK19.18 with and without Dox against MIA-PaCa2 GFP-Nluc CD19+. **a** Representative dot plots of iTRUCK19.18 cell lysis without (left) and with (50 ng/ml) (right) Dox from the first encounter. **b** Representative dot plots of iTRUCK19.18 cell phenotype without (left) and with (50 ng/ml) (right) Dox. **c** Representative histograms of PD1 and TIM3 expression in iTRUCK19.18 cells without (left) and with (50 ng/ml) (right) Dox.

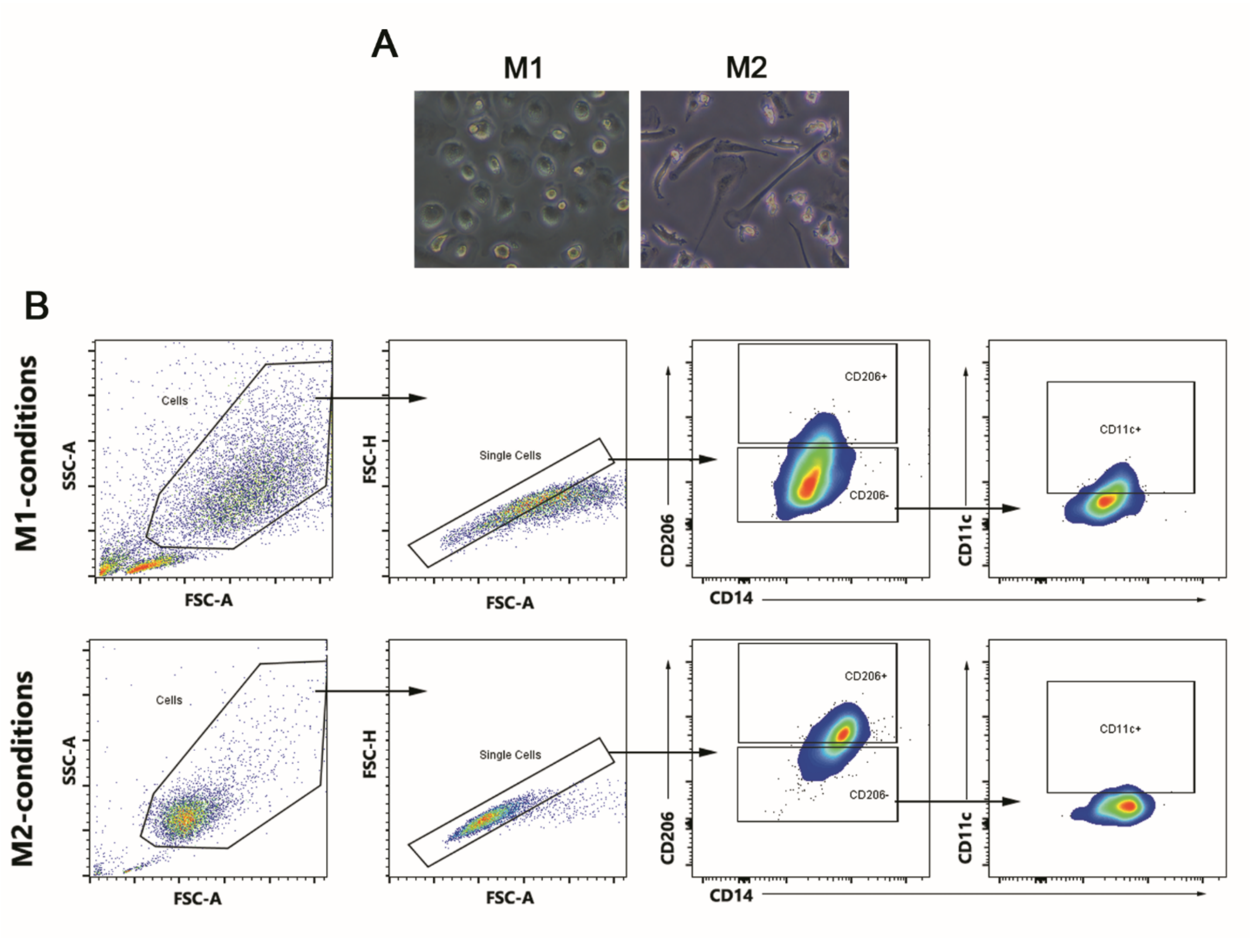


**Fig. S6** Characterization of primary M1 and M2 macrophages. **a** Bright-field microscopy images of human macrophages polarized to M1 (left) or M2 (right) phenotype. **b** Gating strategy for the identification of M1 and M2 macrophages by FACS according to CD14, CD206 and CD11c expression.
